# STK11 is required for the normal program of ciliated cell differentiation in airways

**DOI:** 10.1101/652107

**Authors:** Qiqi Chu, Changfu Yao, Xiangbing Qi, Barry Raymond Stripp, Nan Tang

## Abstract

The functional properties of mucosal surfaces are dependent on establishing the correct proportions of specialized epithelial cell types. Multiciliated cells (also known as ciliated cells) are evolutionarily conserved and functionally indispensable epithelial cells, as suggested by the link between ciliated cell dysfunction and chronic human disease. Ciliated cell differentiation is an ordered process that involves initial cell fate determination and multiciliogenesis. STK11, a serine/threonine kinase, has been reported to be down-regulated in human diseases associated with ciliopathies and functions as a tumor suppressor. Here, we show that STK11 is a physiological factor for the normal program of ciliated cell differentiation by phosphorylating MARK3, which directly suppresses ERK1/2 mediated pRB inactivation. Loss of *Stk11* in airway progenitors impairs the differentiation of ciliated cells in both embryonic and adult airways. Our study establishes that STK11/MARK3/ERK1/2 signaling cascade is a key regulator to integrate ciliated cell fate commitment and the subsequent process of multiciliogenesis.

## Introduction

The airway epithelium serves critical functions to protect the lung from environmental insults. Two major types of epithelial cells in the airway epithelium, secretory and multiciliated cells (also known as ciliated cells), provide airway secretions and ensure removal of invading pathogens and inhaled particles^1^. It is known that the balance between secretory and ciliated cells is fundamental for normal airway function. Impaired differentiation or abnormal functions of secretory and/or ciliated cells lead to many respiratory diseases, including airway infections, cystic fibrosis lung disease, and chronic obstruction pulmonary disease^1^. Therefore, the differentiation of secretory and ciliated cells needs to be tightly regulated during development, homeostasis, and regeneration.

Ciliated cells are derived from multipotent endodermal progenitor cells of the embryonic lung and either secretory or basal progenitors of the postnatal airway^2^. Seminal work has demonstrated that Notch signaling, an evolutionarily conserved cell fate determinant, plays a key role in controlling the balance between secretory versus ciliated cell fates in both embryonic and adult lungs^3-10^. Activation of Notch signaling results in more secretory cells^4,6,8,10^, while inhibition results in an increased number of ciliated cells and a decreased number of secretory cells^3,4,6,9,11^. It is known that in response to cell growth and cytokine signaling, Notch also works with a significantly more complex regulatory network, including p53 and STAT3 signaling, to control epithelial cell differentiation^11,12^. In addition to the function of these signaling networks in ciliated cell fate specification, multiple genes, such as Multicillin and E2F4, have been shown to regulate a subsequent multiciliogenesis process by inducing expression of centriole biogenesis genes and promoting centriole assembly^13-15^. However, the mechanisms that integrate these factors to coordinate the step-by-step process of ciliated cell differentiation remain largely unknown.

Here, using single-cell RNA-sequencing analysis to characterize ciliated cell differentiation in adult airways, we found that the tumor suppressor gene *Stk11* is simultaneously induced at the time of ciliated cell fate determination. STK11, a serine/threonine kinase, has been reported to be down-regulated in human diseases associated with ciliopathies, including Bardet–Biedl syndrome (BBS)^16^, Polycystic kidney disease (PKD)^17^ and Nephronophthisis (NPHP)^18^. We found here that *Stk11* was highly enriched in ciliated cells in both embryonic and adult lungs, and deleting *Stk11* specifically from epithelial progenitor cells impairs ciliated cell differentiation in both embryonic and adult lungs. We have demonstrated that a STK11/MARK3/ERK1/2 signaling cascade acts to enforce ciliated cell lineage commitment and multiciliogenesis in embryonic and adult airways.

## Results

### The expression of *Stk11* is associated with the differentiation of ciliated cells in the airway epithelium

To characterize the differentiation program of ciliated cells, we isolated EpCAM^+^CD45^-^ CD31^-^ intropulmonary epithelial cells from the adult mouse lungs and performed single-cell RNA-Sequencing (scRNA-seq) analysis. In total, 5,842 epithelial cells were used for integrating scRNA-seq analysis. 6 cell clusters were identified by the expression of known marker genes, including alveolar type II cells (AT2); alveolar type I cells (AT1); proximal club cells (Prox-Club)^19^; distal club cells (Dis-Club)^19^; cells that express both *Scgb1a1* and *Foxj1* (Club/Ciliated); and ciliated cells (Ciliated) (Supplementary Fig. S1a).

To better characterize the differentiation program of ciliated cells, we extracted 1,767 cells in Prox-Club, Dis-Club, Club/Ciliated, and Cililated clusters and performed further analyses (Fig. 1a). Interestingly, we found that *Stk11*, which was well known for its functions as a tumor suppressor and frequently mutated in non-small cell lung cancer, was abundantly expressed in cells within the Club/Ciliated and Ciliated cell clusters^20^ (Fig. 1a, Supplementary Fig. S1b). We further described the trajectory of ciliated cell differentiation using a diffusion map and characterized the pseudo-temporal expression levels of genes during ciliated cell differentiation (Fig. 1b). We found that the expression level of *Stk11* simultaneously increased during ciliated cell differentiation process (Fig. 1b).

**Fig. 1.**
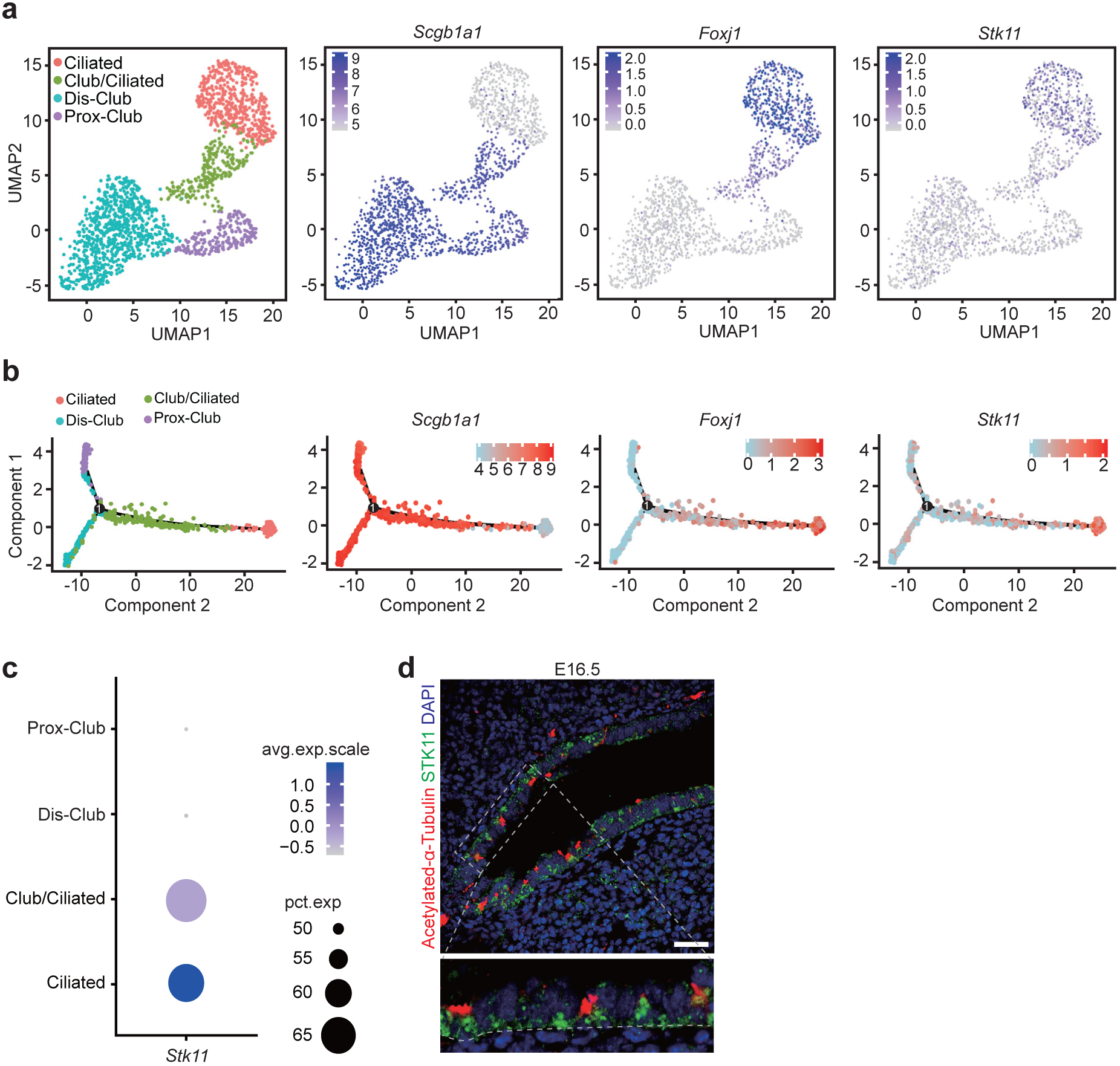
The expression level of *Stk11* is associated with the differentiation of ciliated cells in airways. **a** The UMAP plots of scRNA-seq analyses show the expression of *Scgb1a1*, *Foxj1*, or *Stk11* in adult airway epithelial cells from adult lungs. **b** The pseudo-time trajectories show that the expression level of *Scgb1a1* decreased during the ciliated cell differentiation process, whereas the expression levels of *Foxj1* and *Stk11* increased during the ciliated cell differentiation process. **c** The dot plot of *Stk11* expression score in different cell types. The dot size represents the proportion of cells in a cluster that express the gene. The dot color corresponds to the average expression level of the gene. **d** E16.5 lungs were stained with antibodies against Acetylated-α-Tubulin and STK11. Scale bars: 25 μm.

The decreased expression level of Stk11 has been reported in human ciliopathies^16-18^. By calculating the enrichment score of *Stk11* in different cell clusters, we found that *Stk11* is highly enriched in cells of both Club/Ciliated and Ciliated clusters in adult lungs (Fig. 1c). By immunofluorescence staining with antibodies against Acetylated-α-tubulin and STK11, we found that STK11 was expressed among many airway epithelial cells of embryonic day (E) 16.5 lungs, including a majority of Acetylated-α-tubulin^+^ ciliated cells and some undifferentiated cells, but rarely expressed in SCGB1A1^+^ cells (Fig. 1d, Supplementary Fig. S1c). Together, these results suggest that Stk11 may function in regulating the differentiation of ciliated cells.

### Deletion of *Stk11* in endodermal progenitors of embryonic lung or secretory progenitors of adult lung impairs ciliated cell differentiation

To explore the potential roles of STK11 in ciliated cell differentiation of the developing airway, *Shh*-Cre; *Stk11*^F/F^ mice (*Shh*-*Stk11*) were generated, in which *Stk11* is specifically deleted in endodermal progenitors of the embryonic lung epithelium. In the developing mouse airway, ciliated cells are generated from endodermal progenitor cells. MYB^+^ ciliated cells appear between E13.5 and E14.5, and express the transcription factor FOXJ1^+^ as they differentiate into mature ciliated cells^21^. Epithelial cell differentiation was evaluated in lungs of E16.5 *Shh*-*Stk11* and *Shh*-Cre; *Stk11*^F/+^ (Control) mice by immunofluorescence. The abundance of FOXJ1^+^ and MYB^+^ ciliated cells were each significantly decreased, whereas the percentage of SCGB1A1^+^ club cells was significantly increased, in both tracheal and intrapulmonary epithelia of E16.5 *Shh*-*Stk11* mice, as compared to their littermate Controls (Fig. 2a, b, Supplementary Fig. S2a-d). Notably, excluding the possibility that the impaired ciliated cell differentiation in the *Shh*-*Stk11* lungs results from a developmental delay, we observed similar changes in E18.5 and E20.5 *Shh*-*Stk11* lungs as compared to Control lungs (Supplementary Fig. S2e-h).

**Fig. 2.**
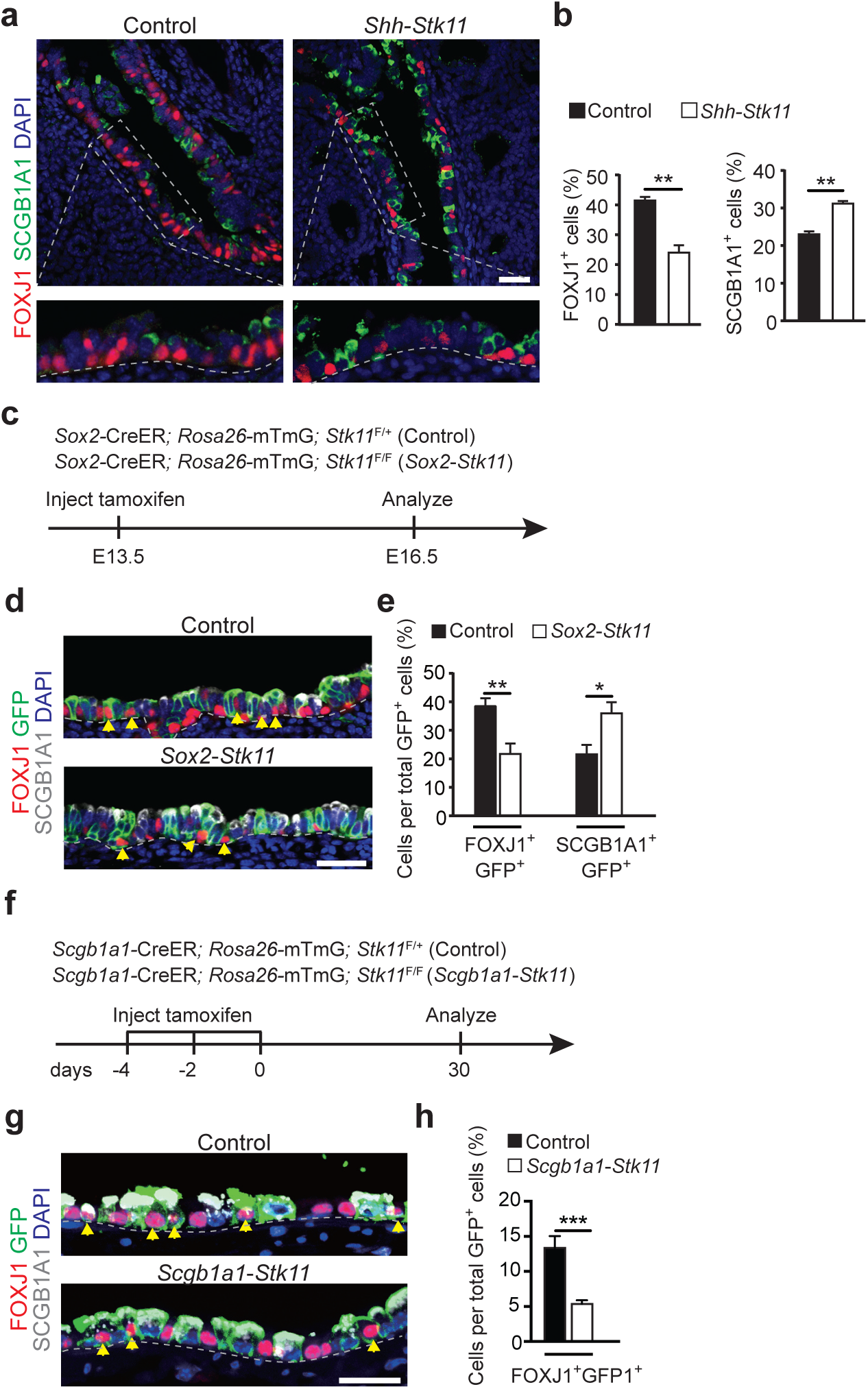
Loss of *Stk11* in airway progenitor cells impairs ciliated cell differentiation. **a** E16.5 lungs were stained with antibodies against FOXJ1 and SCGB1A1. **b** The proportion of FOXJ1^+^ cells (Left, n=3) and SCGB1A1^+^ cells (Right, n=4) in the intrapulmonary airways of E16.5 lungs. **c** Pregnant mice carrying *Sox2*-CreER; *Rosa26*-mTmG; *Stk11*^F/+^ (Control) and *Sox2*-CreER; *Rosa26*-mTmG; *Stk11*^F/F^ (*Sox2*-*Stk11*) embryos were treated with tamoxifen at E13.5 and analyzed at E16.5. **d** E16.5 lungs were stained with antibodies against GFP, FOXJ1, and SCGB1A1. The yellow arrowheads indicate cells expressing both GFP and FOXJ1. **e** The proportion of FOXJ1^+^GFP^+^ cells to GFP^+^ cells (Left) (Control, n=6; *Sox2*-*Stk11*, n=5) and SCGB1A1^+^GFP^+^ cells to GFP^+^ cells (Right) (n=3) in the intrapulmonary airways of E16.5 lungs. **f** 10 weeks-old mice were treated with three doses of tamoxifen and analyzed after 30 days. **g** Lungs were stained with antibodies against GFP, FOXJ1, and SCGB1A1 at day 30. The yellow arrowheads indicate cells expressing both GFP and FOXJ1. **h** The proportion of FOXJ1^+^GFP^+^ cells to GFP^+^ cells at day 30 (n=4). *, *P*<0.05; **, *P*<0.01; ***, *P*<0.001. Data shown in the graphs are means ± SEM. Student’s *t*-test. Scale bars: **a**, **d**: 25 μm, **g**: 20 μm.

During embryonic lung development, endodermal progenitors give rise not only to club and ciliated cells but also to basal and neuroendocrine cells. Therefore, we further analyzed the differentiation of basal cells and neuroendocrine cells in the epithelium of E16.5 *Shh*-*Stk11* and Control lungs. The proportion of basal cells (as a function of total tracheal epithelia) was not significantly different between lungs of *Shh*-*Stk11* and Control embryos (Supplementary Fig. S2i, j). In addition, we observed the proportion of neuroendocrine cells was not significantly different in both genotypes of lungs (Supplementary Fig. S2k, l). These results establish that the loss of *Stk11* does not significantly affect the differentiation of basal cells or neuroendocrine cells in embryonic lungs.

We further investigated the effect of conditionally deleting *Stk11* in SOX2^+^ airway progenitor cells. It has been shown that SOX2 is expressed exclusively in airway progenitor cells during embryonic lung development^22^. Pregnant females carrying *Sox2*-CreER; *Stk11*^F/F^; *Rosa26*-mTmG (*Sox2*-*Stk11*) and *Sox2*-CreER; *Stk11*^F/+^; *Rosa26*-mTmG (Control) embryos were injected with tamoxifen at E13.5 (Fig. 2c), a stage at which the SOX2^+^ airway progenitor cells have not started to differentiate into club or ciliated cells. At E16.5, we found that the proportion of FOXJ1^+^ ciliated cells was significantly decreased and that the proportion of SCGB1A1^+^ club cells was significantly increased among lineage-labeled intrapulmonary airway epithelial cells in *Sox2*-*Stk11* lungs as compared to those of Control embryos (Fig. 2d, e).

Club cells in the adult airway function as secretory progenitor cells that can self-renew and slowly differentiate into ciliated cells to maintain the correct proportions of specialized epithelial cell types^12,23,24^. We next sought to investigate the role of STK11 in regulating ciliated cell differentiation in the adult airway. To achieve this, *Scgb1a1*-CreER; *Stk11*^F/F^; *Rosa26*-mTmG (*Scgb1a1*-*Stk11*) adult mice were injected with three doses of tamoxifen for conditional deletion of *Stk11* within airway club cells and were monitored for 30 days to determine the impact on ciliated cell differentiation (Fig. 2f). The percentage of newly differentiated ciliated cells among lineage-labeled cells decreased significantly in *Scgb1a1*-*Stk11* mice (Fig. 2g, h). These results together with those for embryonic lung support a role for STK11 in controlling ciliated cell differentiation.

### The cell–cycle exit and ciliated cell differentiation are controlled by STK11 within embryonic and adult airways

To further explore the mechanisms through which STK11 regulates ciliated cell differentiation, RNA-seq was performed to compare the transcriptomes of E16.5 *Shh*-*Stk11* and Control mice. Consistent with our immunostaining results, we found that ciliated cell markers (*e.g.*, *Mcidas*, *Myb*, and *Foxj1*^25^) were significantly reduced and found that secretory cell markers, including a club cell marker *Scgb1a1* and a mucous cell marker *Muc5b*, were significantly increased in the lungs of *Shh*-*Stk11* embryos (Fig. 3a-c, Supplementary Dataset S1). We performed gene ontology (GO) analysis to identify the enriched terms amongst the differentially expressed genes in *Shh*-*Stk11* lungs and also found that many of the significantly upregulated genes have known roles as positive regulators of cell proliferation and cell-cycle progression (*e.g.*, *Ranbp2*^26^, *Smc5*^27^, *Mki67*^28^, *Sgo2a*^29^, *Taf1*^30^, *Ccnd2*^31^, and *Cenpe*^32^ (Fig. 3a, b, Supplementary Fig. S3a, and Supplementary Dataset S1). Many of the significantly downregulated genes are known to participate in cilium movement and assembly (Fig. 3a, c, Supplementary Fig. S3b, and Supplementary Dataset S1). These results indicate that both ciliated cell differentiation and cell-cycle status are significantly affected in the *Shh*-*Stk11* lungs.

**Fig. 3.**
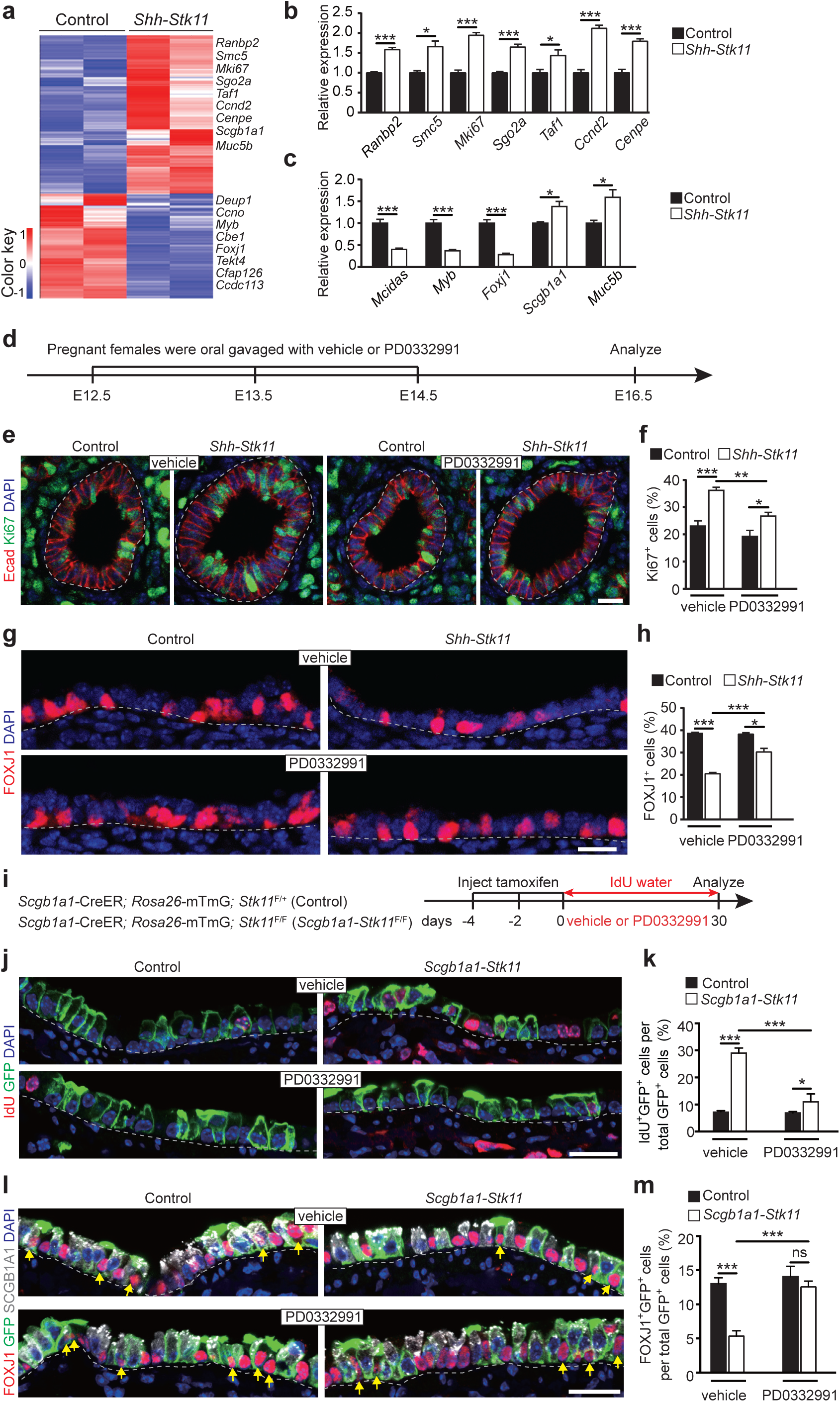
Inhibiting cell proliferation partially rescues the impaired ciliated cell differentiation. **a** Heat map of the expression values of genes expressed differentially between E16.5 *Shh*-*Stk11* lungs and control lungs. **b** Fold change of relative mRNA expression of airway epithelial cell genes in E16.5 *Shh*-*Stk11* lungs compared to Control lungs (n=3). **c** Fold change of relative mRNA expression of cell-cycle genes in E16.5 *Shh*-*Stk11* lungs compared to Control lungs (n=3). **d** Pregnant females were oral gavaged daily with a CDK4/6 inhibitor (PD0332991) each day from E12.5 to E14.5. Lungs were analyzed at E16.5. **e** Cell proliferation was analyzed by antibody staining against Ki67 and E-cadherin (E-cad). **f** The proportion of Ki67^+^ cells in the intrapulmonary airways of E16.5 lungs (vehicle Control, n=5; vehicle *Shh*-*Stk11*, n=3; PD0332991 Control, n=7; PD0332991 *Shh*-*Stk11*, n=8). **g** Immunofluorescence staining with antibodies against FOXJ1 and SCGB1A1 in lungs after either vehicle or PD0332991 treatment. **h** The proportion of FOXJ1^+^ cells (vehicle Control, n=3; vehicle *Shh*-*Stk11*, n=4; PD0332991 Control, n=3; PD0332991 *Shh*-*Stk11*, n=5) and the proportion of SCGB1A1^+^ cells (n=3) in the intrapulmonary airways of E16.5 lungs after either vehicle or PD0332991 treatment. **i** 10 week-old mice were treated with three doses of tamoxifen. Mice were treated with PD0332991 or vehicle every other day via oral gavage and fed with water containing IdU for 30 days before collecting lungs for analysis. **j** Immunofluorescence staining with antibodies against IdU and GFP in lungs after either vehicle or PD0332991 treatment for 30 days. **k** The proportion of IdU^+^ cells in the GFP labeled intrapulmonary airway epithelium (n=3). **l** Immunofluorescence staining with antibodies against FOXJ1, GFP, and SCGB1A1 in lungs after either vehicle or PD0332991 treatment for 30 days. The yellow arrows indicate cells expressing both GFP and FOXJ1. **m** The proportion of FOXJ1^+^GFP^+^ cells to GFP^+^ cells in lungs after either vehicle or PD0332991 treatment for 30 days (n=3). *, *P*<0.05; **, *P*<0.01; ***, *P*<0.001. Data shown in the graphs are means ± SEM. Student’s *t*-test. Scale bars: **e**, **g**: 11 μm; **j**, **l**: 20 μm.

We therefore analyzed the cell-cycle status of airway epithelial cells at E16.5 by propidium iodide (PI) staining, and found that 15.4% of the EpCAM^+^ cells had entered the G2/M phase in *Shh*-*Stk11* lungs, whereas only 11% of the EpCAM^+^ cells entered the G2/M phase in Control lungs (Supplementary Fig. S3c). Consistent with this PI staining result, the proportions of PH3^+^ (phospho-histone H3, a mitosis-specific marker) and Ki67^+^ intrapulmonary airway epithelial cells (Ki67 is a protein present during all active phases of the cell cycle; *i.e.*, G_1_, S, G_2_, and mitosis) were significantly increased in *Shh*-*Stk11* lungs compared to Control lungs (Supplementary Fig. S3d-g). In addition, loss of *Stk11* does not induce apoptosis in embryonic lungs (Supplementary Fig. S3h). We also investigated the impact of STK11 on proliferation of epithelial cells in adult airways. *Scgb1a1*-*Stk11* and Control mice were given IdU water for 30 days prior to sacrifice, and the cumulative IdU-labeling index was determined for lineage-labeled intrapulmonary airway epithelial cells (Supplementary Fig. S3i): there were significantly more IdU^+^ lineage-labeled epithelial cells in *Scgb1a1*-*Stk11* mice than that in Control mice (Supplementary Fig. S3j, k).

### Inhibiting cell proliferation partially rescues the defective ciliated cell differentiation in airways of *Stk11*-deficient mice

Considering that STK11 is known to suppress tumor growth by inhibiting cell proliferation^33,34^, we hypothesize that deletion of *Stk11* either in endodermal progenitors of developing lung or in secretory progenitors of adult lung, prevents these progenitor cells from exiting an active cycling state that is necessary for ciliated cell differentiation. To test this, we determined whether inhibitors of the cell cycle could rescue the ciliated cell differentiation defect in lungs of *Shh*-*Stk11* and/or *Scgb1a1*-*Stk11* mice. Pregnant females carrying *Shh*-*Stk11* and Control embryos were treated with PD0332991, a CDK4/6 inhibitor^35,36^, or vehicle once daily via oral gavage from E12.5 to E14.5, and embryonic lungs collected at E16.5 for analysis (Fig. 3d). CDK4/6 are known to initiate the phosphorylation of pRB, which disrupts its binding with E2Fs and allows cell-cycle entry^37^. We confirmed that the PD0332991 treatment could specifically reduce the number of p-pRB^+^ cells in both the Control and *Shh*-*Stk11* lungs (Supplementary Fig. S3l, m). Interestingly, we found that Control lungs had a much lower proportion of p-pRB^+^Ki67^+^ epithelial cells than did *Shh*-*Stk11* lungs, indicating that the cell proliferation in the *Stk11*-wild type epithelium is largely driven by p-pRB-independent regulatory mechanisms. Loss of *Stk11* only activates p-pRB-dependent cell-cycle progression (Supplementary Fig. S3n, o), demonstrating that STK11-dependent and STK11-independent regulation simultaneously occur in airway progenitor cells. Not surprisingly, the proliferative response of the intrapulmonary airway epithelium of *Shh*-*Stk11* lungs was significantly attenuated by the PD0332991 treatment (Fig. 3e, f). Importantly, the proportion of ciliated cells in intrapulmonary airways of PD0332991-treated *Shh*-*Stk11* embryos was significantly restored compared to those of vehicle-treated *Shh*-*Stk11* embryos (Fig. 3g, h).

Next, we treated *Scgb1a1*-*Stk11* mice with PD0332991 or vehicle once every other day for 30 days (Fig. 3i). PD0332991 treatment significantly decreased the proportion of IdU^+^ lineage traced cells in *Scgb1a1*-*Stk11* mice (Fig. 3j, k). Decreased proliferation in airways of PD0332991-treated *Scgb1a1*-*Stk11* mice restored ciliated cell differentiation to levels observed in vehicle-treated Control intrapulmonary airways (Fig. 3l, m). Taken together, these results indicate that the inhibition of cell-cycle progression in secretory progenitor cells of adult airways, a process regulated by STK11, allows the differentiation of ciliated cells.

### Ciliated cell differentiation requires the kinase activity of STK11

STK11 is a serine/threonine protein kinase that controls the activity of multiple downstream kinases which fulfill diverse roles in embryonic development and adult tissue homeostasis^38-40^. To understand the role of STK11 kinase activity in regulating ciliated cell differentiation, we generated recombinant adenoviral vectors expressing either wild type STK11 (Ad-STK11^WT^) or kinase-dead STK11 (Ad-STK11^KD^). The active site lysine residue K78 contributes to both the binding and orientation of ATP, the phosphate donor. Note that the Stk11^K78I^ (STK11^KD^) variant lacks STK11 kinase activity^38^.

*In vitro* trachea cultures were established by (i) removing the tracheal rudiments from E13.5 *Shh*-*Stk11* or littermate Control embryos, (ii) making a longitudinal incision to expose the luminal surface and (iii) culturing lumen-side-up at the air-liquid interface on TransWell permeable membrane supports (Fig. 4a). At E13.5, a stage before basal-luminal segregation^41^, no ciliated cells were found in the tracheal epithelium (Supplementary Fig. S4a). Tracheal rudiments were infected with Ad-GFP, Ad-STK11^WT^, or Ad-STK11^KD^ viruses and cultured for 5 days (Fig. 4a): we found that the proportions of Ki67^+^ epithelial cells were significantly decreased in the *Shh*-*Stk11* tracheal rudiments with Ad-STK11^WT^ infection at post-culture day 5 (Fig. 4b, c). No significant change of proliferation were observed on *Shh*-*Stk11* tracheal rudiments with either Ad-GFP or Ad-STK11^KD^ infection. Interestingly, when we examined Control tracheal rudiments infected with the Ad-STK11^KD^ virus, we observed a modest but significant increase in epithelial cell proliferation (Fig. 4b, c), possibly due to a dominant negative effect of the kinase-dead STK11 mutant. Notably, in all of the cultured tracheal rudiments, very few basal cells were observed at the various developmental stages.

**Fig. 4.**
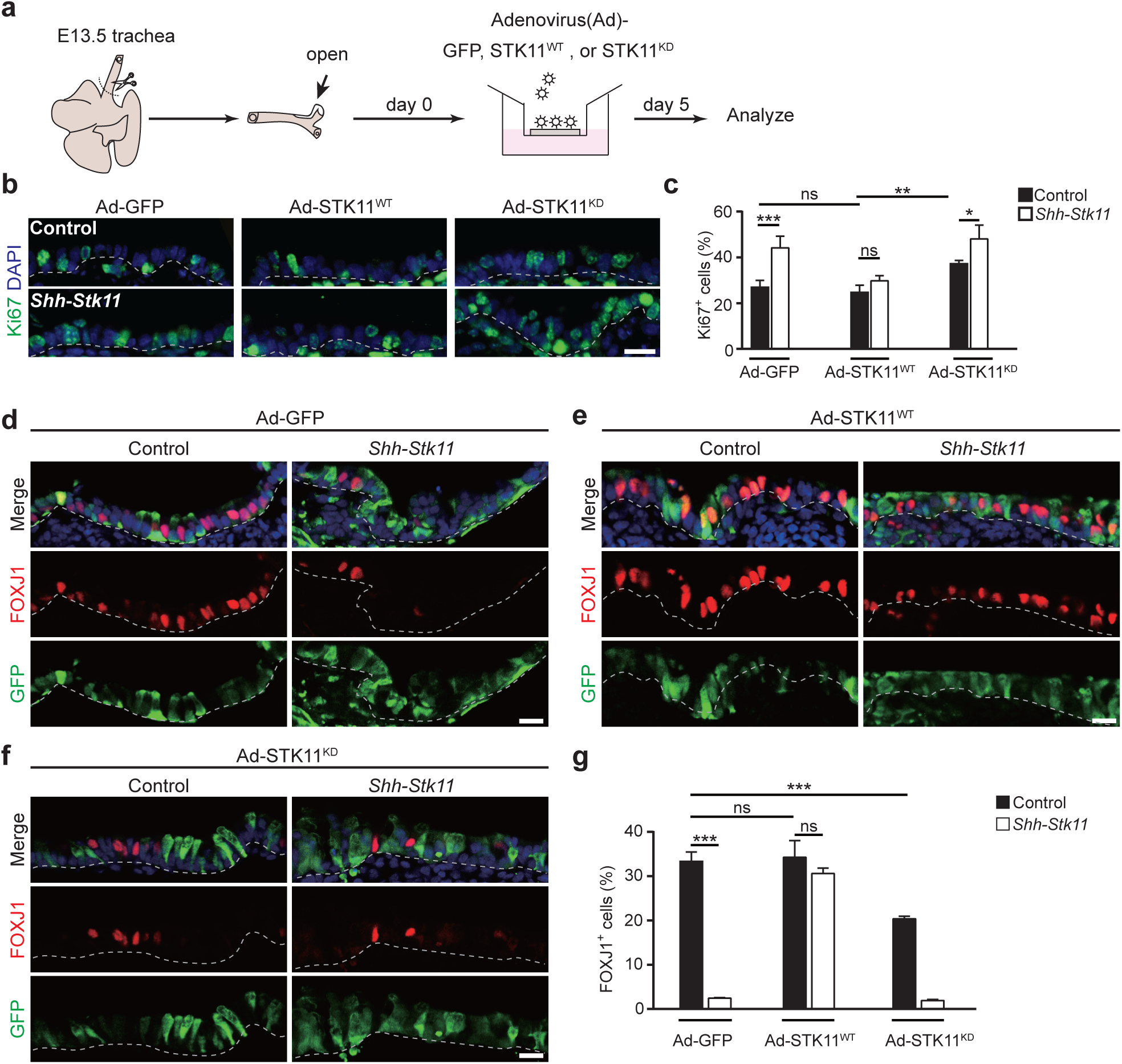
The kinase activity of STK11 is required for ciliated cell differentiation. **a** A schematic illustration showing the air-liquid interphase trachea culture system of this study. Adenovirus (Ad)-GFP, STK11^WT^, or STK11^KD^ particles were added into the medium at day 0. The tracheal rudiments were cultured with different adenovirus for 5 days. **b**-**c** The proportion of Ki67^+^ cells in the epithelium of cultured tracheal rudiments at day 5 (c) were quantified by Ki67 immunostaining (b) (Ad-GFP Control, n=5; Ad-GFP *Shh*-*Stk11*, n=6; Ad-STK11^WT^ Control, n=4; Ad-STK11^WT^ *Shh*-*Stk11*, n=5; Ad-STK11^KD^ Control, n=4; Ad-STK11^KD^ *Shh*-*Stk11*, n=6). **d**-**f** Cultured tracheal rudiments were stained with antibodies against FOXJ1 and GFP at day 5. **g** The proportion of FOXJ1^+^ cells in the epithelium of cultured tracheal rudiments at day 5 (Ad-GFP Control, n=3; Ad-GFP *Shh*-*Stk11*, n=3; Ad-STK11^WT^ Control, n=3; Ad-STK11^WT^ *Shh*-*Stk11*, n=3; Ad-STK11^KD^ Control, n=5; Ad-STK11^KD^ *Shh*-*Stk11*, n=3). ns, not significant; *, *P*<0.05; **, *P*<0.01; ***, *P*<0.001. Data shown in the graphs are means ± SEM. Student’s *t*-test. Scale bars: 11 μm.

We also analyzed ciliated cell formation in cultured tracheal rudiments by performing immunofluorescence staining with an antibody against FOXJ1. The lack of ciliated cell differentiation in *Shh*-*Stk11* tracheal rudiments was rescued by STK11^WT^, but not by the kinase dead STK11^KD^ (Fig. 4d-g). Consistent with the Ki67 immunostaining results showing a dominant negative effect of STK11^KD^ expression in Control tracheal rudiments, the percentage of ciliated cells in the Control tracheal rudiments infected by the Ad-STK11^KD^ virus decreased significantly compared with those infected by the Ad-GFP virus (Fig. 4d, f, g).

We also conducted experiments in which we re-expressed STK11 within *Shh*-*Stk11* and Control tracheal rudiments. Interestingly, this re-expression of STK11 within *Shh*-*Stk11* tracheal rudiments restored the normal proportion of differentiated ciliated cells without inducing ectopic ciliated cell formation (Fig. 4e, g). Additionally, PD0332991 treatment significantly increased the proportion of ciliated cells in the *Shh*-*Stk11* tracheal rudiments but failed to induce ectopic ciliated cell formation in the Control tracheal rudiments (Supplementary Fig. S4b-d). These results indicate that STK11 is necessary but not sufficient for ciliated cell differentiation. Viewed together, these results demonstrate that the kinase activity of STK11 is essential for the function of STK11 in regulating ciliated cell differentiation.

### Phosphorylation of MARK3 by STK11 is required for ciliated cell differentiation

We next investigated downstream phosphorylation targets of STK11^39^. qPCR analysis showed that two of 13 targets are expressed at substantial levels in the airway epithelium of E16.5 lungs: protein kinase AMP-activated catalytic alpha1 subunit (PRKAA1, also known as AMPKα1) and microtubule affinity-regulating kinase 3(MARK3) (Fig. 5a, Supplementary Fig. S5a, b). Both PRKAA1 and MARK3 are regulated by STK11-mediated phosphorylation^39^. To determine the relative activity of PRKAA1 and MARK3 within developing mouse lung, we performed immunostaining of embryonic lung tissue with phospho-status-specific antibodies. We found that many ciliated cells present in E16.5 lungs had phospho-MARK3 expression (Fig. 5a). Furthermore, the phosphorylation levels of both PRKAA1 and MARK3 were significantly decreased in epithelial cells of *Shh*-*Stk11* embryos compared to the Control, confirming that STK11 is necessary for their activation (Fig. 5a, Supplementary Fig. S5b). We also found, both *in vivo* and *in vitro*, that adenosine monophosphate 5-Aminoimidazole-4-carboxamide ribonucleotide (AICAR), an activator of PRKAA1^42,43^, had no impact on either the proliferative status of airway epithelial cells or ciliated cell differentiation (Supplementary Fig. S5c-l); we there therefore subsequently focused on roles for MARK3 as a downstream mediator of STK11.

**Fig. 5.**
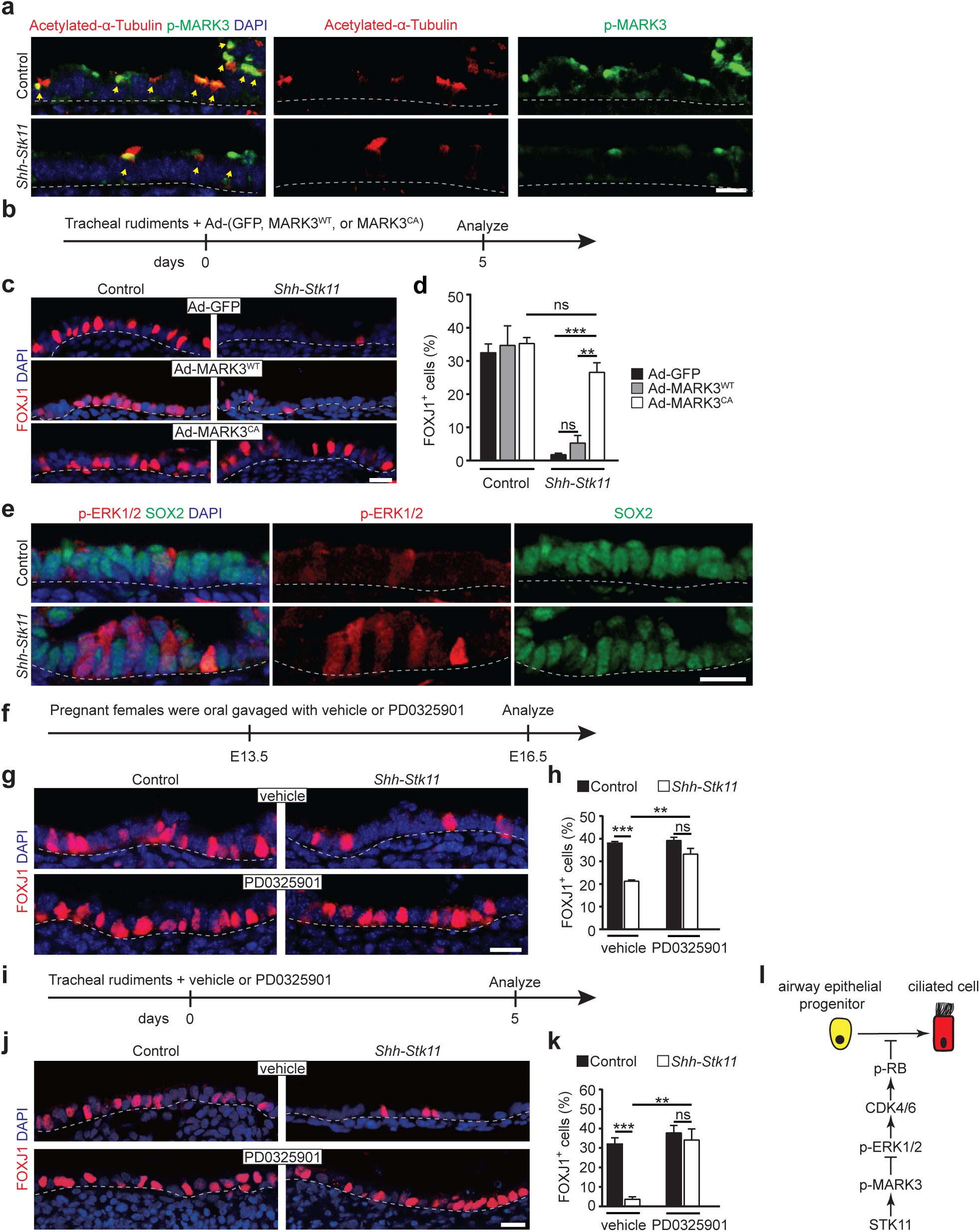
A STK11/MARK3 cascade controls ciliated cell differentiation by limiting the activity of ERK1/2 signaling. **a** Immunofluorescence staining with antibodies against p-MARK3 and Acetylated-α-Tubulin in E16.5 lungs. Yellow arrows indicate ciliated cells expressing both Acetylated-α-Tubulin and p-MARK3. **b** E13.5 tracheal rudiments were cultured with Ad-GFP, Ad-MARK3^WT^, or Ad-MARK3^CA^ viruses for 5 days. **c**-**d** Cultured tracheal rudiments were stained with antibodies against FOXJ1 at day 5 (c). The proportion of FOXJ1^+^ ciliated cells were quantified (Ad-GFP Control, n=3; Ad-GFP *Shh*-*Stk11*, n=3; Ad-MARK3^WT^ Control, n=3; Ad-MARK3^WT^ *Shh*-*Stk11*, n=4; Ad-MARK3^CA^ Control, n=4; Ad-MARK3^CA^ *Shh*-*Stk11*, n=3) (d). **e** E16.5 lungs were stained with antibodies against p-ERK1/2 and SOX2. **f** Pregnant female mice were oral gavaged with p-ERK1/2 inhibitor PD0325901 or vehicle at E13.5. Lungs were analyzed at E16.5. **g**-**h** Immunofluorescence staining with antibodies against FOXJ1 in the vehicle-treated or PD0325901 treated lungs (g). The proportion of FOXJ1^+^ cells (n=3) and the proportion of SCGB1A1^+^ cells (n=3) in the intrapulmonary airways of E16.5 lungs were quantified (h). **i** E13.5 tracheal rudiments were cultured with vehicle or PD0325901 for 5 days. **j**-**k** Cultured tracheal rudiments were stained with antibodies against FOXJ1 at day 5 (j). The proportion of FOXJ1^+^ ciliated cells were quantified (vehicle Control, n=4; vehicle *Shh*-*Stk11*, n=3; PD0325901 Control, n=3; PD0325901 *Shh*-*Stk11*, n=5) (k). **l** A schematic illustration showing that STK11-mediated signaling cascade accounts for the normal program of ciliated cell differentiation in airways. ns, not significant; *, *P*<0.05; **, *P*<0.01; ***, *P*<0.001. Data shown in the graphs are means ± SEM. Student’s *t*-test. Scale bars: 11 μm.

We used recombinant adenoviral vectors expressing a Flag-tagged constitutively activated MARK3 mutant (Ad-MARK3^CA^, the substitution of threonine 211 by a glutamic acid) to examine its role as a downstream target of STK11 that may mediates its effects on ciliated cell differentiation. Tracheal rudiment cultures prepared from E13.5 *Shh*-*Stk11* or Control embryos were infected with Ad-GFP, Ad-MARK3^WT^, or Ad-MARK3^CA^ viruses, followed by culturing for 5 days (Fig. 5b, Supplementary Fig. S6a). Infection of *Shh*-*Stk11* tracheal rudiments with the Ad-MARK3^CA^ virus led to a significant reduction in the proportion of Ki67^+^ epithelial cells compared to tracheal rudiments infected with either the Ad-GFP or Ad-MARK3^WT^ virus (Supplementary Fig. S6b, c). These results show that the proliferative defect observed in rudiments with STK11 deletion can be rescued by the activation of MARK3; such activation actually reduced the proliferative index of the *Shh*-*Stk11* epithelium to the level seen in the Control epithelium.

Examination of the ciliated cells in these virus-infected tracheal rudiments further supported a direct role for activated MARK3 in ciliated cell differentiation. Specifically, infection of *Shh*-*Stk11* tracheal rudiments with the Ad-MARK3^CA^ virus significantly increased the percentage of ciliated cells as compared to *Shh*-*Stk11* tracheal rudiments infected with the Ad-GFP or Ad-MARK3^WT^ viruses (Fig. 5c, d). Thus, our support that STK11-dependent phosphorylation of MARK3 is required for cell-cycle exit and ciliated cell differentiation during airway epithelial development.

### A STK11/MARK3 signaling cascade controls ciliated cell differentiation by limiting ERK1/2 signaling activity

Given the known function of MARK3 as an inhibitor of ERK1/2 signaling^44,45^—a pro-mitotic signaling pathway^46,47^—we next determined whether the effects of p-MARK3 on cell-cycle progression and ciliated cell differentiation within airway epithelial progenitor cells are mediated through inhibition of ERK1/2. To test this, we initially analyzed the presence of p-ERK1/2 in SOX2^+^ intrapulmonary airway epithelial cells of E16.5 Control and *Shh*-*Stk11* lungs. SOX2^+^ epithelial cells of *Shh*-*Stk11* lungs showed a significant increase in p-ERK1/2 immunoreactivity compared to epithelial cells of Control lungs (Fig. 5e). Moreover, the increased expression level of p-ERK1/2 was observed in cultured *Shh*-*Stk11* tracheal rudiments, and expression of MARK3^CA^ in cultured *Shh*-*Stk11* tracheal rudiments significantly decreased the number of epithelial cells showing p-ERK1/2 immunoreactivity (Supplementary Fig. S6d), findings that support a mechanism wherein p-MARK3 controls cell-cycle exit via inhibition of ERK1/2 signaling.

To examine the role of ERK1/2 signaling in ciliated cell differentiation, we treated pregnant females with a specific MEK1/2 inhibitor (PD0325901) or vehicle control at E13.5 and analyzed lungs at E16.5 (Fig. 5f). The number of cells expressing p-ERK1/2 in PD0325901-treated *Shh*-*Stk11* lungs was significantly decreased compared with vehicle-treated *Shh*-*Stk11* lungs (Supplementary Fig. S6e). Both quantification of Ki67^+^ intrapulmonary airway epithelial cells and PI staining showed that the proportion of proliferating epithelial cells in *Shh*-*Stk11* lungs was significantly reduced by the PD0325901 treatment (Supplementary Fig. S6f, g). Importantly, these experiments also showed that PD0325901 treatment rescued the impaired ciliated cell differentiation in the lungs of *Shh*-*Stk11* embryos *in vivo* (Fig. 5g, h) and in the cultured *Shh*-*Stk11* tracheal rudiments *in vitro* (Fig. 5i-k). Moreover, these results show that increased ERK1/2 signaling in *Shh*-*Stk11* lungs promotes cell-cycle progression and interferes with the differentiation of ciliated cells. Thus, our findings define a STK11/MARK3/ERK1/2 signaling cascade allows a normal differentiation program of ciliated cells (Fig. 5l).

### A STK11/MARK3/ERK1/2 signaling cascade functions to enforce ciliated cell lineage commitment

Considering our results demonstrating that the requirement of STK11/MARK3/ERK1/2 signaling cascade in regulating ciliated cell differentiation, we were interested in understanding the relationship between STK11 and ciliated cell fate specification. It is firmly established that Notch signaling determines the fate of lung airway epithelial cells by promoting secretory over ciliated cell fates^4,6,8,10^. We therefore compared the mRNA expression levels of multiple Notch ligands, receptors, and downstream targets in both Control and *Shh*-*Stk11* lungs of E13.5 embryos, a stage at which airway epithelial progenitor cells have not committed to either secretory or ciliated cell fates (Supplementary Fig. S7a). The similar expression levels of Notch pathway genes (*Notch1*, *Notch2*, *Jag1*, *Hes1*, and *Hey1*) that we observed in the lungs of *Shh*-*Stk11* embryos clearly implied that STK11 does not directly regulate the expression of canonical Notch pathway genes.

To further investigate the possible relationship(s) between STK11 and Notch signaling, we generated *Shh*-Cre; *Stk11*^F/F^; *Rbpj*^F/F^ mice (*Shh*-*Stk11*; *Rbpj*) mice. Inactivation of Notch the signaling pathway by deleting *Rbpj*, a DNA-binding protein that associates with NICD (Notch intracellular domain) to transcriptionally regulate downstream genes expression^48^, promotes ciliated cell development and when Notch signaling is absent almost all airway epithelial progenitor cells differentiate into ciliated cells^4,6,8^. We first analyzed the cell proliferation in lungs of E13.5 embryos and found the proportions of PH3^+^ airway epithelial cells were not significantly different between Control and *Shh*-Cre; *Rbpj*^F/F^ (*Shh*-*Rbpj*) lungs. The increased proportions of PH3^+^ epithelial cells were not significantly different between the *Shh-Stk11* and the double mutant lungs (Supplementary Fig. S7b, c). This result is consistent with a previous study which found that the loss of *Rbpj* does not influence cell proliferation in the adult airway epithelium^10^. Additionally, this finding further confirms our conclusion that STK11-regulated cell-cycle exit functions independently of Notch signaling activity.

Having established the independence of STK11-regulated cell-cycle exit and Notch signaling, we were interested to determine whether these pathways exert their effects on ciliated cell differentiation simultaneously. We therefore assessed the proportions of club cells vs. ciliated cells in the lungs of the aforementioned four genotypes by immunostaining using antibodies against SCGB1A1 and FOXJ1. In E16.5 lungs of *Shh*-*Rbpj* mice, 90.56 ± 0.34% intrapulmonary airway epithelial cells were FOXJ1^+^ ciliated cells, and 5.34 ± 0.13% of the intrapulmonary airway epithelial cells expressed neither SCGB1A1 nor FOXJ1; we did not find detectable apoptosis in these lungs (Supplementary Figs. S3h and 7d). Interestingly, the proportion of FOXJ1^+^ ciliated cells was significantly decreased in E16.5 *Shh*-*Stk11*; *Rbpj* lungs (52.23 ± 1.28%), and the proportion of SCGB1A1^-^FOXJ1^-^ intrapulmonary airway epithelial cells increased to 43.4 ± 1.89% in E16.5 *Shh*-*Stk11*; *Rbpj* lungs (Fig. 6a, b). However, by E18.5, the proportion of FOXJ1^+^ ciliated cells in *Shh*-*Stk11*; *Rbpj* lungs increased to 90.83 ± 1.92% (Fig. 6c, d). In addition, we found that many of these SCGB1A1^-^FOXJ1^-^ cells were positive for Ki67 in E16.5 *Shh-Stk11; Rbpj* lungs (Supplementary Fig. S7e, f). Therefore, these results indicate that the SCGB1A1^-^FOXJ1^-^ cells in E16.5 *Shh*-*Stk11*; *Rbpj* lungs ultimately differentiated into FOXJ1^+^ ciliated cells at E18.5, suggesting that these SCGB1A1^-^ FOXJ1^-^ cells have committed to an early ciliated cell fate (pre-ciliated cells). Thus, as the lack of Notch signaling strongly promoted differentiation of airway epithelial cells toward a ciliated cell fate in *Shh-Stk11; Rbpj* lungs, we can conclude that the regulatory influence of STK11/MARK3/ERK1/2 signaling cascade on ciliated cell differentiation is not related to ciliated cell fate specification.

**Fig. 6.**
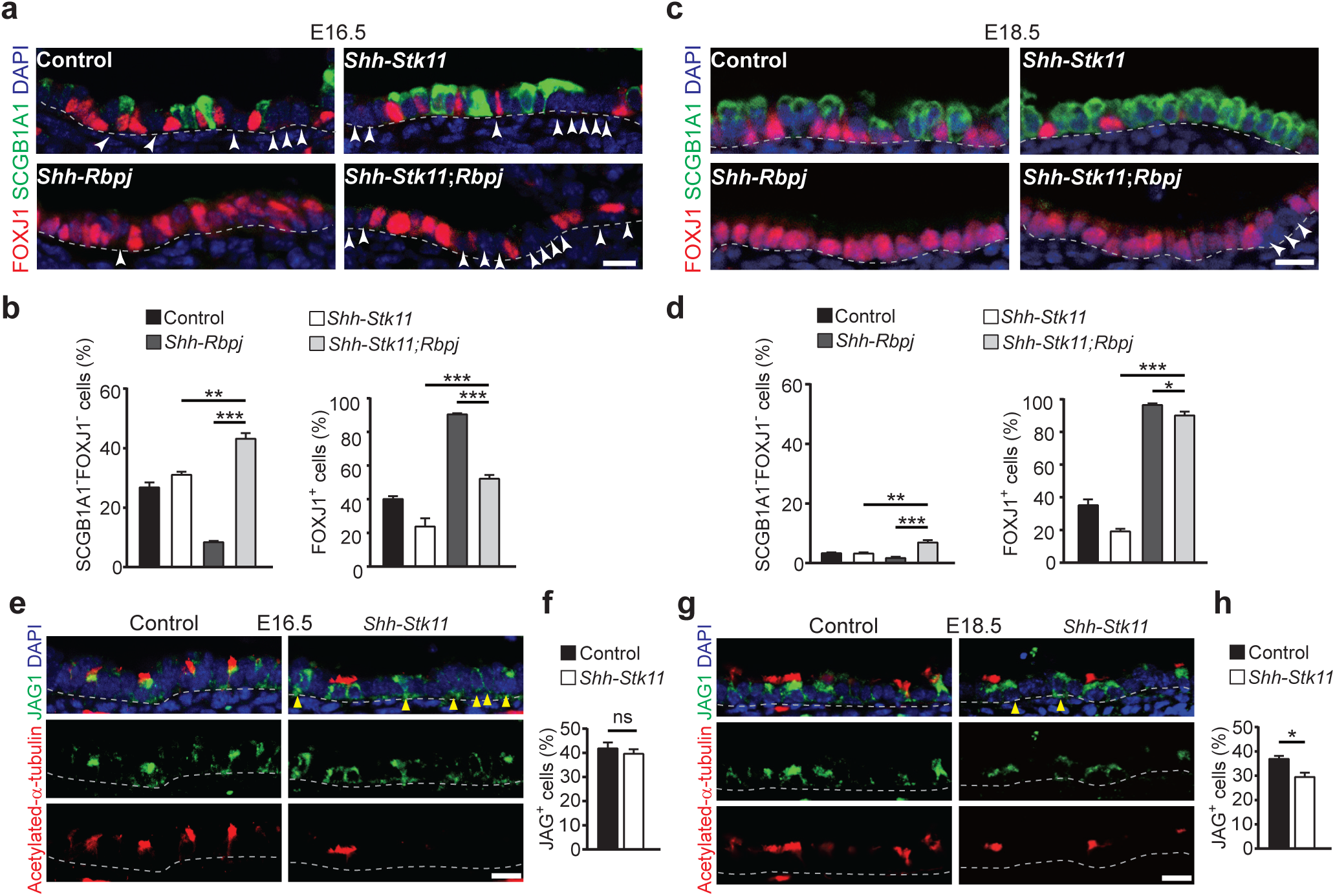
STK11 ensures the ciliated cell fate commitment. **a** Immunofluorescence staining with antibodies against FOXJ1 and SCGB1A1 in the E16.5 lungs. White arrowheads indicate cells that do not express SCGB1A1 and FOXJ1. **b** The proportion of SCGB1A1^-^FOXJ1^-^ cells (Left), and the proportion of FOXJ1^+^ cells (Right) in the intrapulmonary airways of E16.5 lungs (n=3). **c** Immunofluorescence staining with antibodies against FOXJ1 and SCGB1A1 in the E18.5 lungs. White arrowheads indicate cells that do not express SCGB1A1 and FOXJ1. **d** The proportion of SCGB1A1^-^FOXJ1^-^ cells (Left), and the proportion of FOXJ1^+^ cells (Right) in the intrapulmonary airways of E18.5 lungs (n=3). **e** Immunofluorescence staining with antibodies against Acetylated-α-tubulin and JAG1 (JAGGED1) in E16.5 lungs. Yellow arrowheads indicate the Acetylated-α-tubulin^-^ cells that express JAG1. **f** The proportion of JAG1^+^ cells in the intrapulmonary airways of E16.5 lungs (Control, n=4; *Shh*-*Stk11*, n=4). **g** Immunofluorescence staining with antibodies against Acetylated-α-tubulin and JAG1 in E18.5 lungs. Yellow arrowheads indicate the Acetylated-α-tubulin^-^ cells that express JAG1. **h** The proportion of JAG1^+^ cells in the intrapulmonary airways of E18.5 lungs (Control, n=4; *Shh*-*Stk11*, n=5). ns, not significant; *, *P*<0.05; **, *P*<0.01; ***, *P*<0.001. Data shown in the graphs are means ± SEM. Student’s *t*-test. Scale bars: 11 μm.

JAGGED1 (JAG1) is a Notch ligand that is known to be expressed by ciliated cells in the intrapulmonary airways^8^. To investigate the expression level of *Jagged1*, we isolated EpCAM^+^CD45^-^CD31^-^ epithelial cells from Control lungs and *Shh-Stk11* lungs at E14.5, E16.5, and E18.5. Surprisingly, we found that the mRNA level of *Jagged1* was not altered in *Shh-Stk11* lungs at E14.5 and E16.5 (Supplementary Fig. S7g). In light of our previous finding that the proportion of ciliated cells decreased in E16.5 *Shh-Stk11* lungs (Fig. 2a, b), we also performed immunostaining using antibodies against JAGGED1 and Acetylated-α-tubulin. We found that the proportion of JAGGED1^+^ cells was not decreased and that many Acetylated-α-tubulin negative cells expressed JAGGED1 in the E16.5 *Shh*-*Stk11* lungs (Fig. 6e, f). These results suggest that the Acetylated-α-tubulin^-^JAGGED1^+^ cells may have committed to ciliated cell fates but cannot perform multiciliogenesis to terminally differentiate into Acetylated-α-tubulin^+^ ciliated cells in the *Shh*-*Stk11* lungs. At E18.5, both the mRNA level of *Jagged1* and the proportion of JAGGED1^+^ cells were decreased in the *Shh*-*Stk11* lungs, likely due to the significantly reduced proportion of ciliated cells at this stage (Fig. 6g, h, Supplementary Fig. S7g). Taken together, our findings indicate that STK11 acts to ensure ciliated cell fate commitment.

## Discussion

In our study, we identified that STK11—in both embryonic and adult lungs—is required for the normal program of ciliated cell differentiation. In the absence of *Stk11*, ciliated cell differentiation is largely impaired. When we inhibited the phosphorylation level of pRB by the PD0332991 treatment, both the increased cell proliferation and impaired ciliated cell differentiation were restored in *Shh-Stk11* lungs. Furthermore, we demonstrated that STK11 regulates cell-cycle exit and ciliated cell differentiation by phosphorylating its downstream target, MARK3, which in turn leads to the inhibition of ERK1/2 signaling-mediated pRB phosphorylation^37,44,49^. Given the clear evidence that decreased expression of STK11 in human diseases is associated with ciliopathies, we conclude that STK11 is a physiological regulator to enforce ciliated cell fate commitment and multiciliogenesis, ensuring the normal program of ciliated cell differentiation.

Notch signaling controls the balance of ciliated and secretory cell fates in developing airways^8^. We found no evidence to support a direct interaction between the STK11 and Notch signaling pathways. It is established that inactivated Notch signaling induces neuroendocrine cell fate specification^50^. The fact that neuroendocrine cell differentiation is not significantly different between *Shh*-*Stk11* and Control lungs provides further evidence that Notch signaling is not impaired in *Shh*-*Stk11* lungs (Supplementary Fig. S2k, l). The relative expression level of *Jagged1* is not significantly decreased in the E13.5, E14.5, or E16.5 *Shh*-*Stk11* lungs, and the expression of JAGGED1 in many Acetylated-α-tubulin^-^ cells of E16.5 *Shh*-*Stk11* lungs suggest that these airway epithelial progenitors may have acquired the ciliated cell fate.

Once they have adopted a low Notch signaling-dependent ciliated cell fate, the pre-ciliated cells initiate the expression of ciliary genes to launch the multiciliogenesis process. It has been demonstrated that multiciliogenesis is a precise process sequentially regulated by the nucleocytoplasmic translocation of E2F4^13^. Once the ciliated cell fate is determined, nuclear E2F4-DP1-Multicilin complex initiates a transcriptional program of centriole biogenesis and turns on genes that are required for multiciliogenesis (*e.g.*, *Deup1, Myb, Ccno, Foxj1)*^15,21,51-54^. Then, E2F4 translocates to cytoplasm and organizes deuterosome assembly centers to initiate centriole amplification^13^. In our study, we found that both the transcriptional coactivator Multicilin (*Mcidas*) and genes that are transcriptionally regulated by E2F4-DP1-Multicilin complex (*Deup1, Myb, Ccno,* and *Foxj1*), were significantly downregulated in *Shh*-*Stk11* lungs (Fig. 3a, b, Supplementary Dataset S1). This finding demonstrates that loss of *Stk11* impairs the transcriptional program of centriole biogenesis. Furthermore, the increased proportion of club cells observed in *Shh*-*Stk11* lungs suggests that diminished expression of centriole biogenesis genes may causes loss of ciliated cell identity in pre-ciliated cells, allowing these cells to assume a non-ciliated cell fate. We therefore propose that STK11/MARK3/ERK1/2 signaling cascade is required for preventing the plasticity of pre-ciliated cells and ensuring their commitment to multiciliogenesis.

Multiciliated cells are the main constituents of mucociliary epithelia, fulfilling critical roles in epithelia of respiratory, nervous, and reproductive systems by providing the mechanical force to mobilize fluids and external particles across the epithelium. These functions rely on proper ciliated cell differentiation during both development and homeostasis. Understanding the molecular mechanisms underlying the step-by-step ciliated cell differentiation process will provide insight for our future understanding of ciliated cell-related diseases.

## Materials and Methods

### Mice

*Shh*-Cre^55^,*Sox2*-CreER^55^, *Rosa26*-mTmG^56^, *Scgb1a1*-CreER^24^, floxed *Rbpj*^57^, and floxed *Stk11*^58^ mice have been described previously. Mice had mixed genetic backgrounds. Noon of the day on which a vaginal plug was detected was considered E0.5. Tamoxifen (75 μg/g body weight) or corn oil was injected into *Scgb1a1*-CreER, floxed *Stk11, Rosa26*-mTmG mice, and mice carrying *Sox2*-CreER, floxed *Stk11* and their respective control embryos. Mice were killed using CO_2_. All experiments were performed in accordance with the recommendations in the Guide for Care and Use of Laboratory Animals of the National Institute of Biological Sciences, Beijing and Cedars-Sinai Medical Center, Los Angeles. All experiments were approved by the Institutional Review Board at the National Institute of Biological Science, Beijing and Cedars-Sinai Medical Center, Los Angeles.

### Single-cell RNA-seq analysis

CD45^-^CD31^-^EpCAM^+^ lung epithelial cells from embryonic lungs at E16.5 and adult lung of 4 male and 4 female mice (2 batches) were FACS sorted and performed for Single-Cell RNA-seq analysis using the 10x Genomic Chromium system, respectively. CellRanger v2.2.0 software was used with the default settings for de-multiplexing, aligning reads with STAR software to mm10 mouse reference genome, and counting unique molecular identifiers (UMIs). Downstream analysis were performed using Seurat v2.3 R package. UMAP (Uniform Manifold Approximation and Projection) method has been used for visualization of unsupervised clustering. The cell type of each cluster is determined by known markers of individual cell types. Airway cells are further subset for expression and pseudo-time analysis. Pseudo-time trajectories are performed using Monocle 2 R package ^59^.

### FACS and cell-cycle analysis

The E16.5 lungs were cut into pieces smaller than 1 mm^3^ after removing the heart. Tissues were then digested using a neutral protease (Worthington Biochemical Corporation) and DNase I (Roche) at 37°C for 20 minutes. The reaction was stopped using DMEM medium supplemented with 10% FBS. The tissue suspension was passed through a 40 μm mesh filter (Thermo Scientific) to remove undigested tissue fragments. The supernatant was centrifuged at 300g for 10 minutes and resuspended with red blood lysis buffer. After centrifuging again, the cells were resuspended with blocking buffer (5% BSA in DPBS) and stained with PE-Cy7 rat anti-CD31 (1:400, 561410, BD biosciences), APC mouse anti-EpCAM (1:400, 17-5791-82, eBioscience), and FITC rat anti-CD45 (1:400, 553080, BD biosciences) for 20 min on ice. DAPI was used to exclude dead cells before sorting. The EpCAM^+^CD45^-^CD31^-^ cells were then sorted via flow cytometry (Aria II, BD Biosciences). The isolated cells were centrifuged at 300g for 10 minutes. Then they were washed with cold DPBS twice. The cells were resuspended and fixed with pre-cold 70% ethanol for 24 hours at 4°C. For cell-cycle analysis, the fixed cells were stained with propidium iodide (P4170, Sigma) supplemented with Rnase A (CW0600S, cwbiotech Corporation) for 15 min at RT. The cell-cycle status was also analyzed by flow cytometry and the data were processed using Flowjo v10 (BD biosciences).

### Administration of inhibitors to mice

The pregnant females were treated with PD0332991 (150 mg/kg, Sigma) by oral gavage from E12.5 to E14.5 each day or with PD0325901 (5 mg/kg, Sigma) by oral gavage at E13.5 or with AICAR (250 mg/kg, Sigma) by intraperitoneal injection from E12.5 to E14.5 each day. 10 weeks old mice were treated with PD0332991 (50 mg/kg, Sigma) every other day for 30 days by oral gavage.

### Trachea culture

The tracheal rudiments were dissected from E13.5 lungs. These tracheal rudiments were made a longitudinal incision with a sharp needle on ice as quickly as possible. The tracheal rudiments were then washed three times in HBSS (Gibco, 14025-092) supplemented with penicillin/streptomycin for 5 minutes on ice. We next transferred these tracheal rudiments to TransWell units and cultured them at an air-liquid interface of DMEM/F12 medium (Gibco, 11039-021). The tracheal rudiments were cultured for 5 days *in vitro*. The day when the tracheal rudiments were dissected and cultured in the medium was defined as day 0. For adenovirus infection experiment, the adenovirus was added into the medium at day 0. The virus titer finally used was 10^8^ (PFU/ml). The tracheal rudiments were cultured with adenovirus for 24 hours. For the inhibitor chemical treatment experiments, AICAR (10 μM), PD0325901 (10 μM), or PD0332991 (5 μM) was added to the medium at day 0 and withdraw after 72 hours treatment, respectively. The medium was changed every day in all experiments.

### RNA sequencing and data processing

Total RNA was extracted from E16.5 lungs using a Direct-zol^TM^ RNA MiniPrep kit (ZYMO RESEARCH, R2050). The sequencing was performed at the Sequencing Center of NIBS with SE50 using a Hiseq-2500 instrument. RNA-seq analyses of E16.5 lungs were performed with at least two independent biological replicates. Enriched GO and KEGG pathway terms for RNA-seq differentially expressed gene sets were identified using DAVID. Visualization of RNA-seq was performed using the Integrative Genomics Viewer.

### Immunohistochemistry

Mouse lungs were fixed in 4% paraformaldehyde at 4°C overnight. For frozen sections, lungs were immersed in sucrose, and then embedded in OCT. For paraffin sections, lungs were dehydrated in a gradient of ethanol, made transparent with xylene, and embedded in wax. Immunofluorescence staining was performed using the following primary antibodies: rabbit anti-STK11 antibody (Cell Signaling, 13031, 1:500), mouse anti-Acetylated-α-Tubulin (Sigma, T6793, 1:2000), mouse anti-FOXJ1 (eBioscience, 14-9965, 1:500), mouse anti-MYB (Santa Cruz, sc-74512, 1:500), rabbit anti-Ki67 (Abcam, ab15580, 1:300), chicken anti-GFP antibody (Abcam, ab13970, 1:1000), rabbit anti-AMPKα (PRKAA1) antibody (Cell Signaling, 2535s, 1:500), rat anti-E-Cadherin (Invitrogen clone ECCD-2, 1:500), rabbit anti-PH3 (Millipore, 06-570, 1:100), rabbit anti-SCGB1A1 (Millipore, 07-623, 1:300), rabbit anti-phosphorylated MARKs antibody (Cell Signaling, 4836, 1:200), rabbit anti-Caspase3 antibody (Cell Signaling, 9664s, 1:300), goat anti-SOX2 antibody (Santa Cruz, sc-17320, 1:100), mouse anti-P63 (Santa Cruz, sc-8431, 1:500), rabbit anti-CGRP antibody (Sigma, C8198, 1:300), mouse anti-Flag antibody (Sigma, A2220, 1:200), rabbit anti-phosphorylated ERK1/2 antibody (Cell Signaling, 4370S, 1:200), rabbit anti-p-pRB antibody (Cell Signaling, 8516, 1:1600), rabbit anti-JAG1 antibody (Cell Signaling, 70109, 1:1000), and mouse anti-IdU antibody (BD Biosciences, 347580, 1:500). The following secondary antibodies were used: Alexa Fluor 488-Donkey anti-goat (Jackson Immuno Research), Alexa Fluor Cy3-Donkey anti-rabbit (Jackson Immuno Research), Alexa Fluor 488-Donkey anti-chicken (Jackson Immuno Research), Alexa Fluor 568-Donkey anti-rat (Jackson Immuno Research), and Alexa Fluor 647-Donkey anti-rat (Jackson Immuno Research), Alexa Fluor 647-Donkey anti-rabbit (ThermoFisher Scientific), Alex Fluor 568-Donkey anti-mouse IgG (H+L) (ThermoFisher Scientific). All of the Alexa Fluor coupled secondary antibodies were used at 1:500 dilutions.

Immunohistochemical staining for STK11, phosphorylated-MARKs, MYB, and FOXJ1 were performed using a TSA Biotin Tyramide Amplification system (SAT700001EA, Perkin Elmer) for amplification. The following secondary antibodies were used: Biotin-SP-conjugated AffiniPure Donkey anti-rabbit (Jackson Immuno Research), Biotinylated horse anti-mouse (Vector Laboratories). The cultured tracheas were collected and fixed in 4% paraformaldehyde at 4°C for 1 hour. Then they were immersed in sucrose and embedded in OCT.

### Microscopy and imaging

Tissue section immunofluorescence staining was imaged using confocal microscopy (Leica Corporation). Cells were counted based on the basis of nuclear staining with DAPI and specific cell markers of the respective cell types. Cells were counted using 40x magnification fields covering the whole left lung. Images were processed and analyzed using ImageJ (NIH), Adobe Photoshop (Adobe), and Adobe Illustrator (Adobe).

### Adenovirus production

The genes encoding the kinase-dead variants of *Stk11* and wild-type *Stk11* were cloned from the plasmids kindly provided by Dr. Hongbin Ji. To disable the kinase activity of STK11, the lysine residue at position 78 was mutated into isoleucine. The *Mark3^CA^* gene was synthesized by Obio Technology (Shanghai). The STK11^WT^ adenovirus and STK11^KD^ adenovirus were constructed by cloning wild-type *Stk11* and kinase-dead variants of *Stk11* into the vector pAdeno-MCMV-EGFP-P2A-MSC-3FLAG. The MARK3^CA^ and MARK3^WT^ adenovirus were constructed by cloning the mutant MARK3 and wild type MARK3 into the vector pAdeno-MCMV-3FLAG-P2A-EGFP. The GFP-adenovirus, STK11^WT^ adenovirus, STK11^KD^ adenovirus, MARK3^WT^ adenovirus, and MARK3^CA^ adenovirus were purchased from Obio Technology (Shanghai).

### Analyze the adenovirus infection efficacy

The infection efficacy of Ad-GFP, Ad-STK11^WT^, and Ad-STK11^KD^ was determined by performing immunofluorescence staining with an antibody against GFP. The infection efficacy of Flag-tagged Ad-GFP, Ad-MARK3^WT^ and Ad-MARK3^CA^ was determined by performing immunofluorescence staining with an antibody against Flag.

### RNA extraction and qPCR

The total RNA of E16.5 lungs was extracted using Trizol reagent (Ambion) and Direct-Zol RNA Miniprep kits (ZYMO research). RNA was reverse transcribed into first-strand complementary DNA using HiScript II Q RT SuperMix (Vazyme). The qPCR analysis was performed on a CFX96 Tough instrument (Bio-Rad) using KAPA SYBR FAST qPCR Master Mix (KAPA). The primers were provided in Supplementary Table S1.

### Statistical analysis

All *in vivo* quantifications are based on the airway epithelial cells of the intrapulmonary airways in embryonic lungs. All quantification data are presented as the mean ± SEM. (as indicated in figure legends). All quantification data presented in the figures were collected from at least three experimental replications per group on different days using different mice. We used two-tailed Student’s *t*-tests to assess differences between means.

### Data availability

The RNA-seq data along with their associated meta data have been deposited in the GEO database under accession code GSE116690 and GSE123838. All relevant data of this study are available from the authors upon reasonable request.

## Acknowledgements

We thank Dr. Hongbin Ji for providing us the plasmids of STK11^WT^ and STK11^KD^. We thank for Dr. Xiangrong Guan for technique support. This work was funded by the National Institutes of Health 1T32HL134637, R01HL135163; California Institution of Regenerative Medicine CIRM LA1-06915. This work was funded by Beijing Major Science and Technology Projects, Z171100000417003.

## Conflict of Interest

The authors declare no competing or financial interests.

## Author Contributions

Q.C. and C.Y handled the following: study design, characterization of animal model, immunohistochemistry data analysis and interpretation, and writing of manuscript. X.Q helped the inhibitor experiment. B.R.S. and N.T. co-designed experiments and co-write the manuscript.

