## Supplemental information for "STK11 is required for the normal program of ciliated cell differentiation in airways"

#### The Supplementary information file includes:

##### Supplementary Figs. S1 to S7.

##### Supplementary Table S1.

The list of primers used for qPCR_related to Material and Methods.

##### Supplementary Dataset S1.

The information of RNA-seq datasets_related to Fig 3.

#### Figure S1

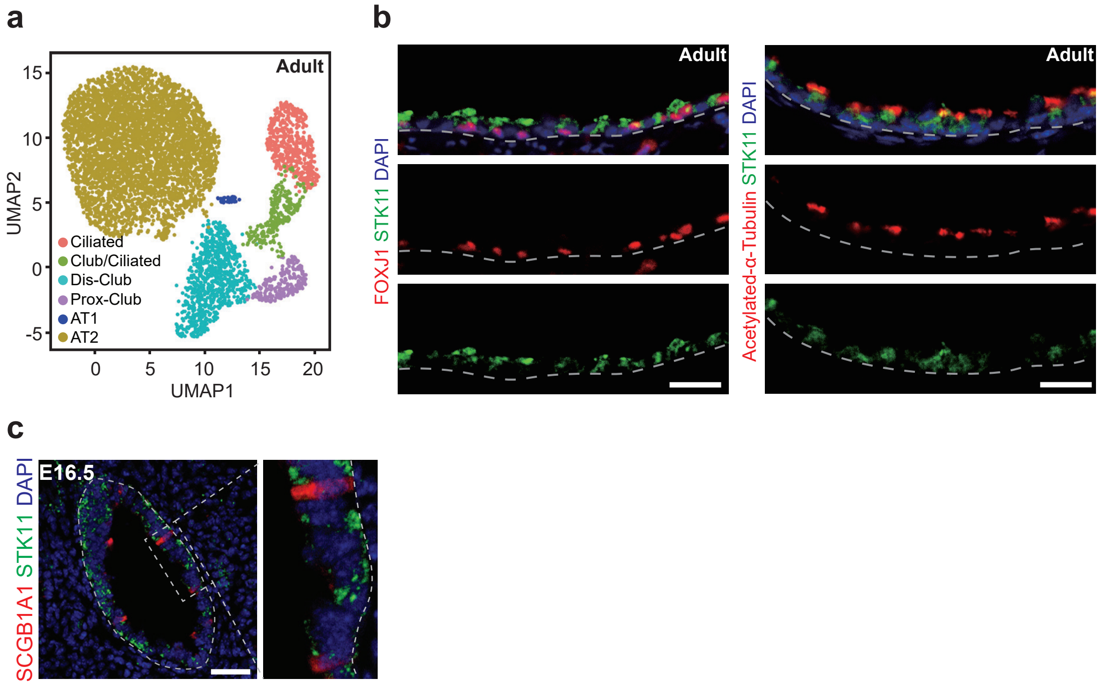
a
Adult
UMAP2
UMAP1
Ciliated
Club/Ciliated
Dis-Club
Prox-Club
AT1
AT2
b
Adult
FOXJ1 STK11 DAPI
Acetylated-α-Tubulin STK11 DAPI
c
E16.5
SCGB1A1 STK11 DAPI

**Figure S1.** Analyzing of *Stk11* during ciliated cell development by scRNA-seq and immunostaining. **a** The UMAP plots of scRNA-seq of epithelial cells from adult lungs showed six clusters. **b** Immunofluorescence staining with antibodies against FOXJ1 (left), Acetylated-a-tubulin (right), and STK11 in lungs of adult mice. **c** Immunofluorescence staining with antibodies against STK11 and SCGB1A1 in E16.5 lungs. Scale bars: **b**: 20 μm; **c**: 25 μm.

### Figure S2

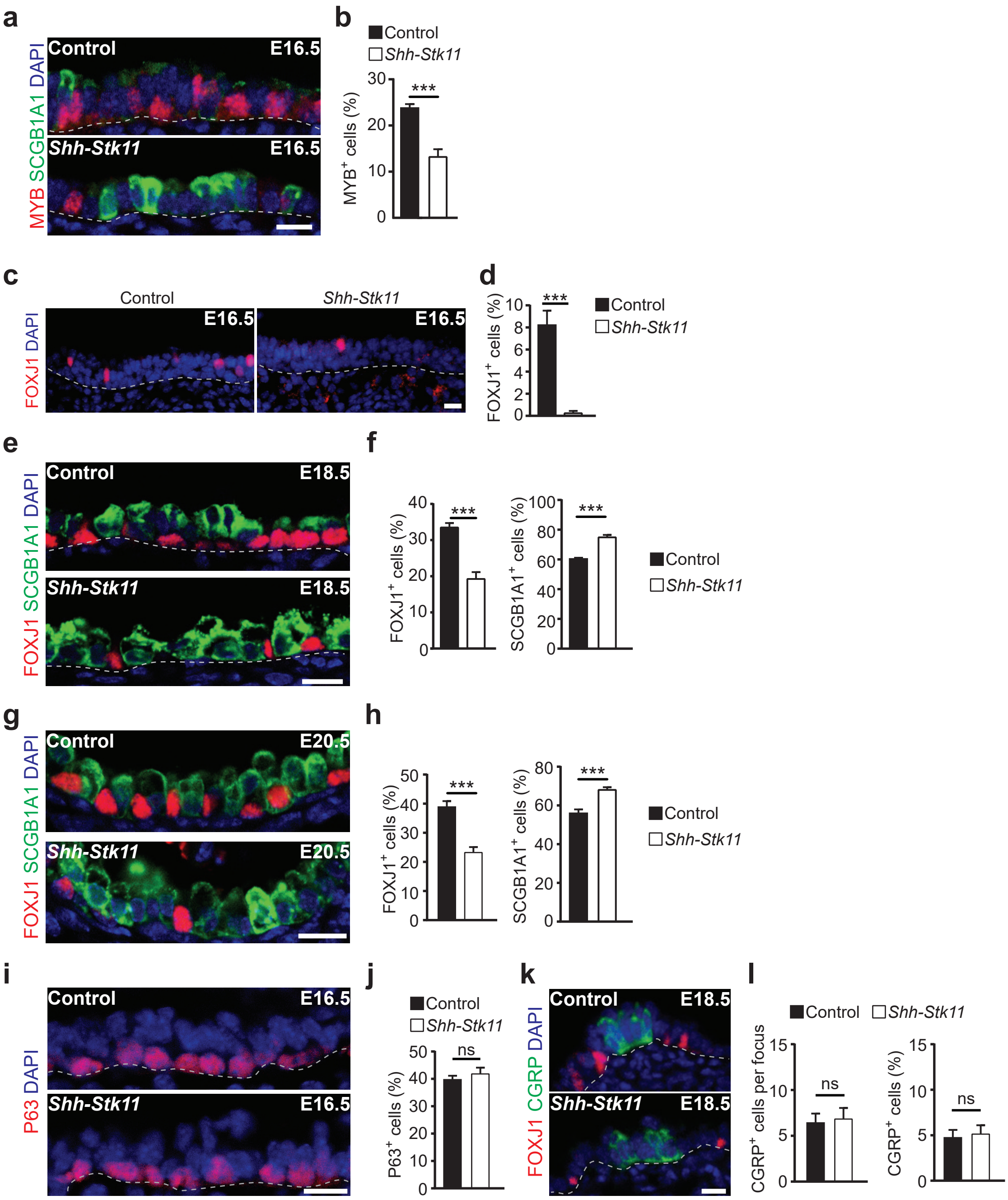

a
Control
E16.5
Shh-Stk11
E16.5
MYB SCGB1A1 DAPI
b
Control
Shh-Stk11
***
MYB+ cells (%)
0
10
20
30
c
Control
Shh-Stk11
E16.5
E16.5
FOXJ1 DAPI
d
***
FOXJ1+ cells (%)
0
2
4
6
8
10
Control
Shh-Stk11
e
Control
E18.5
Shh-Stk11
E18.5
FOXJ1 SCGB1A1 DAPI
f
***
FOXJ1+ cells (%)
SCGB1A1+ cells (%)
Control
Shh-Stk11
g
Control
E20.5
Shh-Stk11
E20.5
FOXJ1 SCGB1A1 DAPI
h
***
FOXJ1+ cells (%)
SCGB1A1+ cells (%)
Control
Shh-Stk11
i
Control
E16.5
Shh-Stk11
E16.5
P63 DAPI
j
Control
Shh-Stk11
ns
P63+ cells (%)
k
Control
E18.5
Shh-Stk11
E18.5
FOXJ1 CGRP DAPI
l
Control
Shh-Stk11
ns
CGRP+ cells per focus
CGRP+ cells (%)

**Figure S2.** Loss of STK11 in airway progenitor cells impairs airway epithelial cell differentiation. **a** Immunofluorescence staining with antibodies against MYB and SCGB1A1 in E16.5 lungs. **b** The proportion of $MYB^{+}$ cells in the intrapulmonary airways of E16.5 lungs (Control, n=3; *Shh-Stk11*$^{F/F}$, n=4). **c** Immunofluorescence staining with an antibody against FOXJ1 in the trachea of E16.5 lungs. **d** The proportion of $FOXJ1^{+}$ ciliated cells in the tracheal epithelium of E16.5 lungs (n=4). **e** Immunofluorescence staining with antibodies against FOXJ1 and SCGB1A1 in E18.5 control lungs and *Shh-Stk11*$^{F/F}$ lungs. **f** The proportion of $FOXJ1^{+}$ cells in E18.5 lungs and the proportion of $SCGB1A1^{+}$ cells in E18.5 lungs (n=3). **g** Immunofluorescence staining with antibodies against FOXJ1 and SCGB1A1 in E20.5 control lungs and *Shh-Stk11*$^{F/F}$ lungs. **h** The proportion of $FOXJ1^{+}$ cells in E20.5 lungs and the proportion of $SCGB1A1^{+}$ cells in E20.5 lungs (Control, n=5; *Shh-Stk11*$^{F/F}$, n=8). **i** Immunofluorescence staining with an antibody against P63 in E16.5 lungs. **j** The proportion of $P63^{+}$ cells in the tracheal epithelium of E16.5 lungs (n=3). **k** Immunofluorescence staining with antibodies against FOXJ1 and CGRP in E18.5 lungs. **l** The $CGRP^{+}$ cell per focus and the proportion of $CGRP^{+}$ cells in E18.5 lungs (n=3). ns, not significant; ***, *P*<0.001. Data shown in the graphs are means ± SEM. Student's *t*-test. Scale bars: 11 μm.

### Figure S3

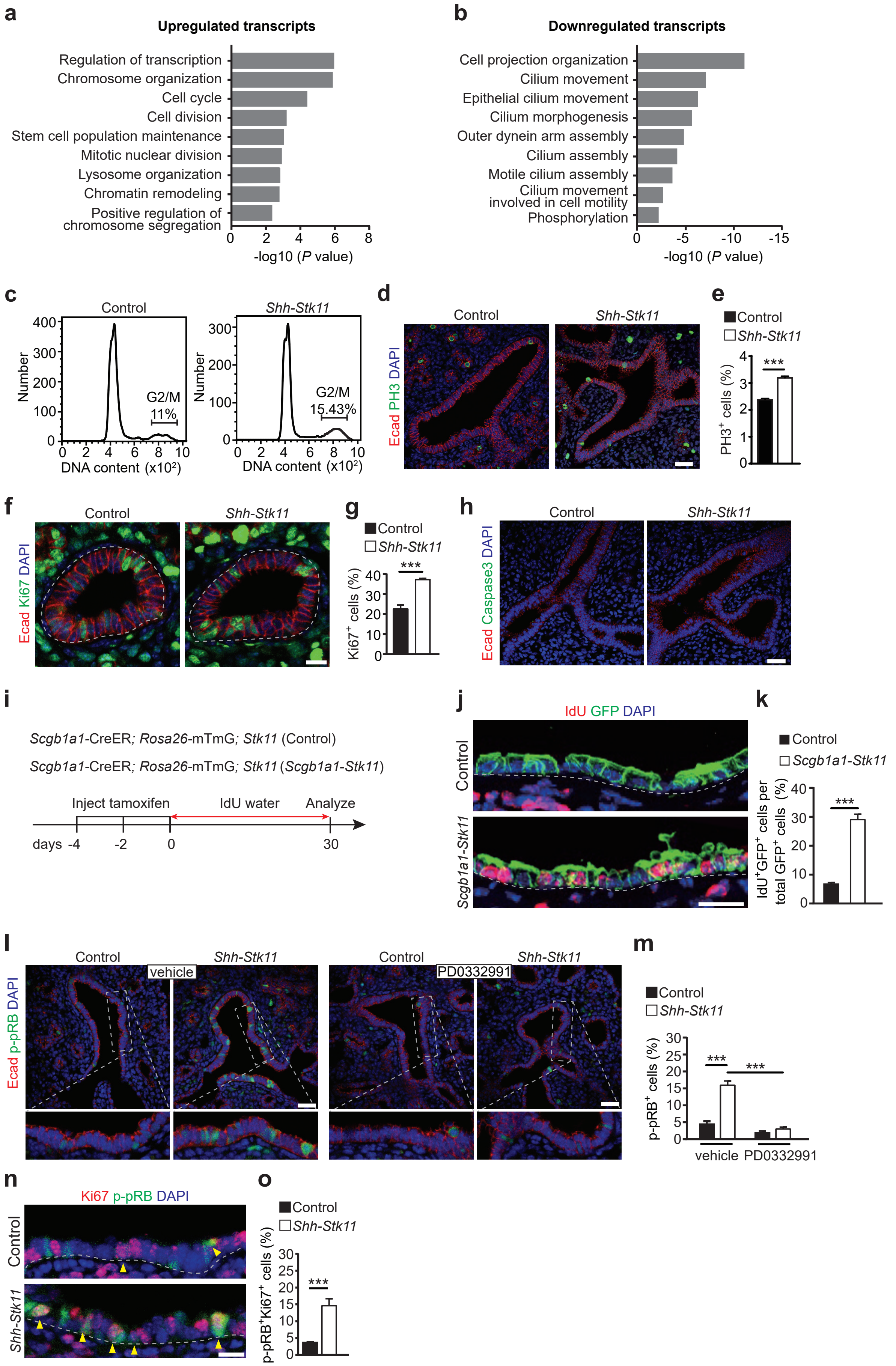

a
Upregulated transcripts
Regulation of transcription
Chromosome organization
Cell cycle
Cell division
Stem cell population maintenance
Mitotic nuclear division
Lysosome organization
Chromatin remodeling
Positive regulation of chromosome segregation
-log10 (P value)
b
Downregulated transcripts
Cell projection organization
Cilium movement
Epithelial cilium movement
Cilium morphogenesis
Outer dynein arm assembly
Cilium assembly
Motile cilium assembly
Cilium movement involved in cell motility
Phosphorylation
-log10 (P value)
c
Control
Shh-Stk11
Number
G2/M 11%
G2/M 15.43%
DNA content (x10²)
d
Control
Shh-Stk11
Ecad PH3 DAPI
e
Control
Shh-Stk11
PH3⁺ cells (%)
***
f
Control
Shh-Stk11
Ecad Ki67 DAPI
g
Control
Shh-Stk11
Ki67⁺ cells (%)
***
h
Control
Shh-Stk11
Ecad Caspase3 DAPI
i
Scgb1a1-CreER; Rosa26-mTmG; Stk11 (Control)
Scgb1a1-CreER; Rosa26-mTmG; Stk11 (Scgb1a1-Stk11)
Inject tamoxifen
IdU water
Analyze
days -4 -2 0 30
j
IdU GFP DAPI
Control
Scgb1a1-Stk11
k
Control
Scgb1a1-Stk11
IdU⁺GFP⁺ cells per total GFP⁺ cells (%)
***
l
Control
Shh-Stk11
vehicle
PD0332991
Ecad p-pRB DAPI
m
Control
Shh-Stk11
p-pRB⁺ cells (%)
***
vehicle
PD0332991
n
Ki67 p-pRB DAPI
Control
Shh-Stk11
o
Control
Shh-Stk11
p-pRB⁺Ki67⁺ cells (%)
***

**Figure S3.** Increased cell proliferation in *Shh-Stk11*$^{F/F}$ lungs. **a-b** DAVID Gene Ontology (GO) analysis of genes that are up-regulated (a) or down-regulated (b) in the E16.5 *Shh-Stk11*$^{F/F}$ lungs as compared to Control lungs. **c** The cell-cycle progression status of epithelial cells in E16.5 lungs was analyzed with PI staining. **d** E16.5 lungs were stained with an antibody against Phospho-Histone H3 (PH3). **e** The proportion of PH3$^{+}$ cells in the intrapulmonary airways of E16.5 lungs (n=3). **f** Immunofluorescence staining with antibodies against Ki67 and Ecad in E16.5 lungs. **g** The proportion of Ki67$^{+}$ intrapulmonary airway epithelial cells in E16.5 lungs (n=4). **h** Immunofluorescence staining with antibodies against Caspase3 and Ecad in E16.5 lungs. **i** 10 week-old *Scgb1a1*-CreER; *Rosa26*-mTmG; *Stk11*$^{F/+}$ (Control) and *Scgb1a1*-CreER; *Rosa26*-mTmG; *Stk11*$^{F/F}$ (*Scgb1a1-Stk11*$^{F/F}$) mice were treated with three doses of tamoxifen and given IdU water for 30 days before analyzing. **j** Immunofluorescence staining with antibodies against IdU and GFP at day 30. **k** The proportion of IdU$^{+}$ cells in the GFP labeled intrapulmonary airway epithelium after IdU water treatment for 30 days (n=4). **l** Immunofluorescence staining with antibodies against Ecad and p-pRB in E16.5 lungs after the vehicle or PD0332991 treatment. **m** The proportion of p-pRB$^{+}$ cells in the intrapulmonary airway epithelium after PD0332991 treatment (vehicle Control, n=3; vehicle *Shh-Stk11*$^{F/F}$, n=3; PD0332991 Control, n=4; PD0332991 *Shh-Stk11*$^{F/F}$, n=3). **n** Immunofluorescence staining with antibodies against Ki67 and p-pRB in E16.5 lungs. **o** The proportion of Ki67$^{+}$p-RB$^{+}$ cells in the Ki67$^{+}$ intrapulmonary airway epithelial cells in E16.5 lungs (n=4). ***, $P<0.001$. Data shown in the graphs are means ± SEM. Student's *t*-test. Scale bars: **d**, **h**, **l**: 25 μm; **f**, **n**: 11 μm. **j**: 20 μm.

### Figure S4

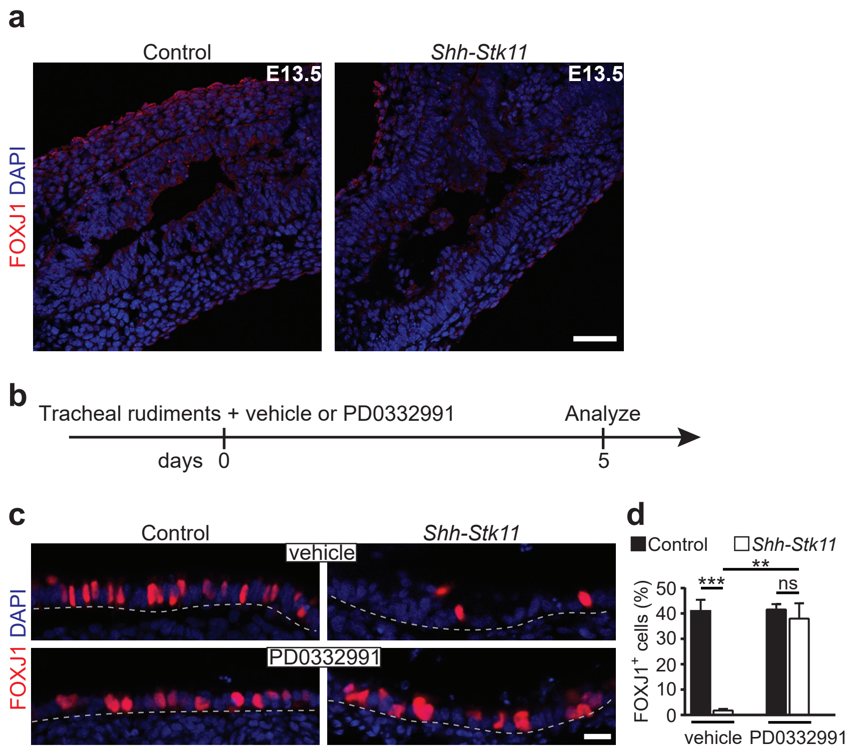
a
Control
Shh-Stk11
E13.5
E13.5
FOXJ1 DAPI
b
Tracheal rudiments + vehicle or PD0332991
Analyze
days 0
5
c
Control
Shh-Stk11
vehicle
PD0332991
FOXJ1 DAPI
d
Control
Shh-Stk11
***
**
ns
FOXJ1+ cells (%)
50
40
30
20
10
0
vehicle
PD0332991

**Figure S4.** Inhibition of cell proliferation can rescue ciliated cell differentiation defect in the *Stk11*-deficient tracheal rudiments. **a** Immunofluorescence staining with an antibody against FOXJ1 in the tracheal rudiments of E13.5 lungs. **b** E13.5 tracheal rudiments were cultured with vehicle or PD0332991 for 5 days. **c** Immunofluorescence staining with an antibody against FOXJ1 in the tracheal rudiments at day 5. **d** The proportion of FOXJ1$^{+}$ ciliated cells in cultured tracheal rudiments at day 5 (n=3). ns, not significant; **, *P*<0.01; ***, *P*<0.001. Data shown in the graphs are means ± SEM. Student's *t*-test. Scale bar: **a**: 25 μm; **c**: 11 μm.

### Figure S5

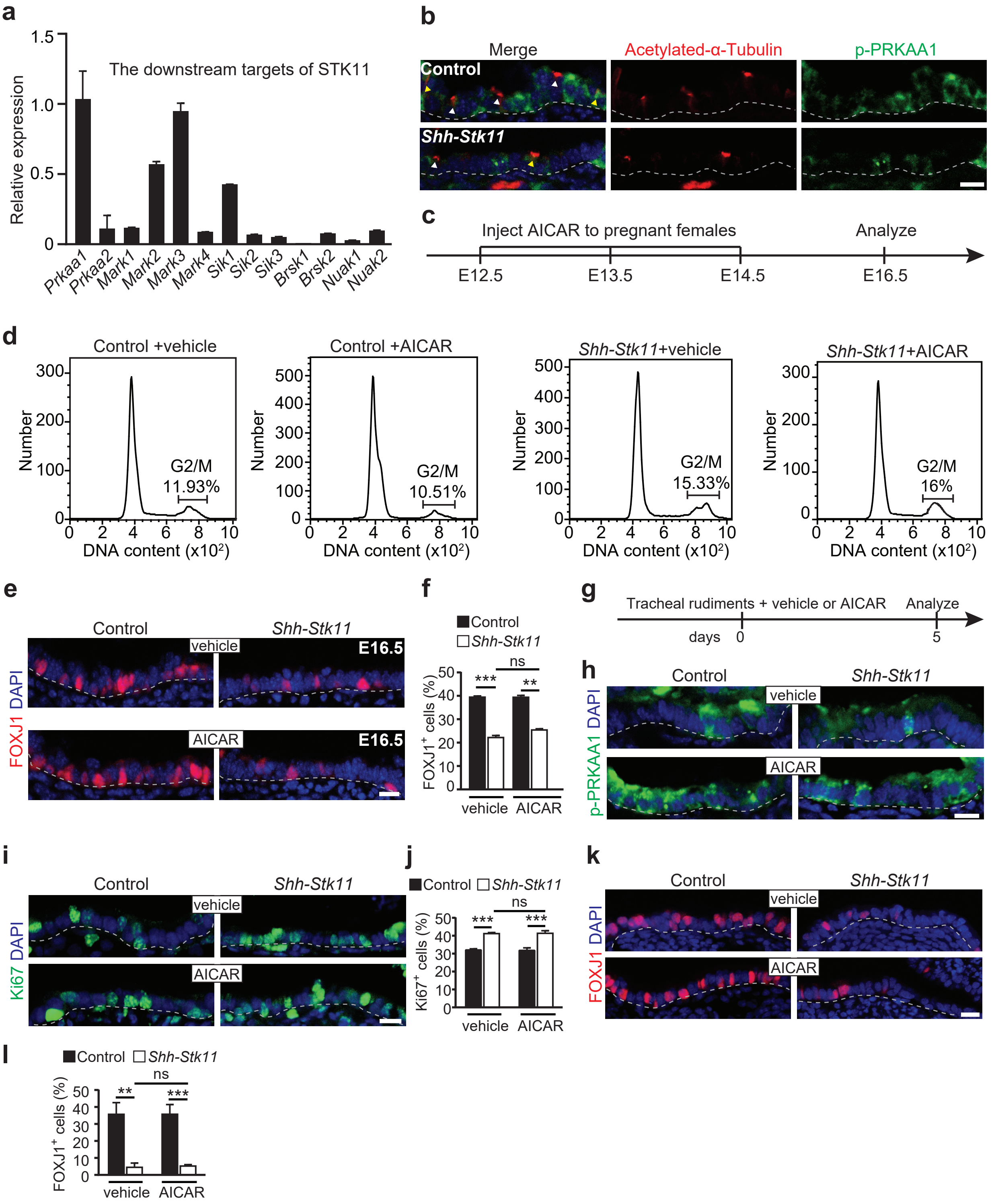

a
The downstream targets of STK11
Relative expression
1.5
1.0
0.5
0
Prkaa1
Prkaa2
Mark1
Mark2
Mark3
Mark4
Sik1
Sik2
Sik3
Brsk1
Brsk2
Nuak1
Nuak2
b
Merge
Acetylated-α-Tubulin
p-PRKAA1
Control
Shh-Stk11
c
Inject AICAR to pregnant females
Analyze
E12.5
E13.5
E14.5
E16.5
d
Control +vehicle
Control +AICAR
Shh-Stk11+vehicle
Shh-Stk11+AICAR
Number
G2/M
11.93%
10.51%
15.33%
16%
DNA content (x10²)
e
Control
Shh-Stk11
vehicle
AICAR
E16.5
FOXJ1 DAPI
f
Control
Shh-Stk11
FOXJ1+ cells (%)
ns
***
**
vehicle
AICAR
g
Tracheal rudiments + vehicle or AICAR
Analyze
days
0
5
h
Control
Shh-Stk11
vehicle
AICAR
p-PRKAA1 DAPI
i
Control
Shh-Stk11
vehicle
AICAR
Ki67 DAPI
j
Control
Shh-Stk11
Ki67+ cells (%)
ns
***
vehicle
AICAR
k
Control
Shh-Stk11
vehicle
AICAR
FOXJ1 DAPI
l
Control
Shh-Stk11
FOXJ1+ cells (%)
ns
**
***
vehicle
AICAR

**Figure S5.** Activating p-PRKAA1 cannot rescue impaired ciliated cell differentiation both *in vivo* and *in vitro*. **a** The mRNA levels of STK11-downstream genes in the epithelial cells of E16.5 lungs (n=3). **b** Immunofluorescence staining with antibodies against p-PRKAA1 and Acetylated-$\alpha$-Tubulin in E16.5 lungs. White arrows indicate ciliated cells positive for Acetylated-$\alpha$-Tubulin but negative for p-PRKAA1. Yellow arrows indicate ciliated cells positive for both Acetylated-$\alpha$-Tubulin and p-PRKAA1. **c** Pregnant mice were treated with AICAR by intraperitoneal injection from E12.5 to E14.5 each day. Lungs were analyzed at E16.5. **d** The cell-cycle progression status of epithelial cells in E16.5 lungs was analyzed by PI staining. **e** Immunofluorescence staining with an antibody against FOXJ1 in the E16.5 lungs after the vehicle or AICAR treatment. **f** The proportion of FOXJ1$^+$ cells in the intrapulmonary airways of E16.5 lungs after the treatments (n=3). **g** E13.5 tracheal rudiments were cultured with AICAR for 5 days. **h** Cultured tracheal rudiments were stained with an antibody against p-PRKAA1 at day 5 after AICAR treatment. **i** Cultured tracheal rudiments were stained with antibodies against Ki67 at day 5. **j** The proportion of Ki67$^+$ epithelial cells in the cultured tracheal rudiments at day 5 (vehicle Control, n=3; vehicle *Shh-Stk11*$^{F/F}$, n=3; AICAR Control, n=3; AICAR *Shh-Stk11*$^{F/F}$, n=4). **k** Cultured tracheal rudiments were stained with an antibody against FOXJ1 at day 5. **l** The proportion of FOXJ1$^+$ ciliated cells in cultured tracheal rudiments at day 5 (vehicle Control, n=4; vehicle *Shh-Stk11*$^{F/F}$, n=4; AICAR Control, n=3; AICAR *Shh-Stk11*$^{F/F}$, n=3). ns, not significant; **, $P$<0.01; ***, $P$<0.001. Data shown in the graphs are means ± SEM. Student's *t*-test. Scale bars: 11 $\mu$m.

### Figure S6

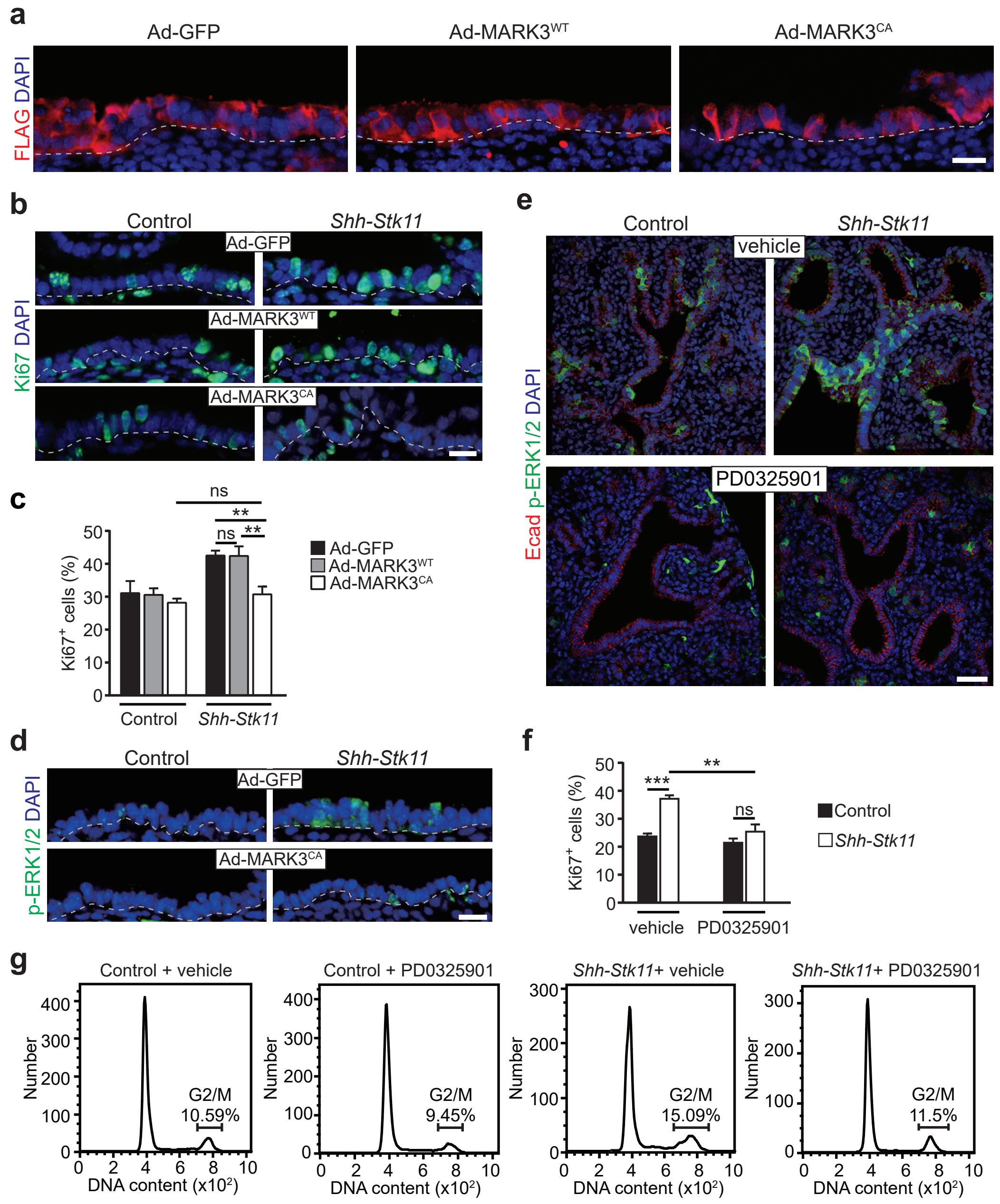

a
Ad-GFP
Ad-MARK3WT
Ad-MARK3CA
FLAG DAPI
b
Control
Shh-Stk11
Ad-GFP
Ad-MARK3WT
Ad-MARK3CA
Ki67 DAPI
c
ns
**
ns
**
Ki67+ cells (%)
Ad-GFP
Ad-MARK3WT
Ad-MARK3CA
Control
Shh-Stk11
d
Control
Shh-Stk11
Ad-GFP
Ad-MARK3CA
p-ERK1/2 DAPI
e
Control
Shh-Stk11
vehicle
PD0325901
Ecad p-ERK1/2 DAPI
f
***
**
ns
Ki67+ cells (%)
Control
Shh-Stk11
vehicle
PD0325901
g
Control + vehicle
Control + PD0325901
Shh-Stk11+ vehicle
Shh-Stk11+ PD0325901
Number
G2/M
10.59%
9.45%
15.09%
11.5%
DNA content (x10²)

**Figure S6.** STK11/MARK3 cascade controls ciliated cell differentiation by limiting the activity of ERK1/2 signaling. **a** Cultured tracheal rudiments were stained with an antibody against Flag at day 5 to indicate the adenovirus infection efficiency. **b** Cultured tracheal rudiments were stained with antibodies against Ki67 at day 5 after adenovirus infection. **c** The proportion of $Ki67^+$ epithelial cells in the cultured tracheal rudiments were quantified (n=3). **d** Cultured tracheal rudiments were stained with antibodies against p-ERK1/2 at day 5 after adenovirus infection. **e** Immunofluorescence staining with antibodies against p-ERK1/2 and Ecad in E16.5 lungs after the vehicle or PD0325901 treatment. **f** The proportion of $Ki67^+$ cells in the intrapulmonary airways of E16.5 lungs after PD0325901 treatment (vehicle Control, n=4; vehicle *Shh*-*Stk11*$^{F/F}$, n=4; PD0325901 Control, n=9; PD0325901 *Shh*-*Stk11*$^{F/F}$, n=7). **g** The cell-cycle status of epithelial cells in E16.5 lungs after the vehicle or PD0325901 treatment were analyzed with PI staining. ns, not significant; **, $P<0.01$; ***, $P<0.001$. Data shown in the graphs are means ± SEM. Student's *t*-test. Scale bars: **a**, **b**, **d**: 11 μm; **e**: 25 μm.

**Figure S7**

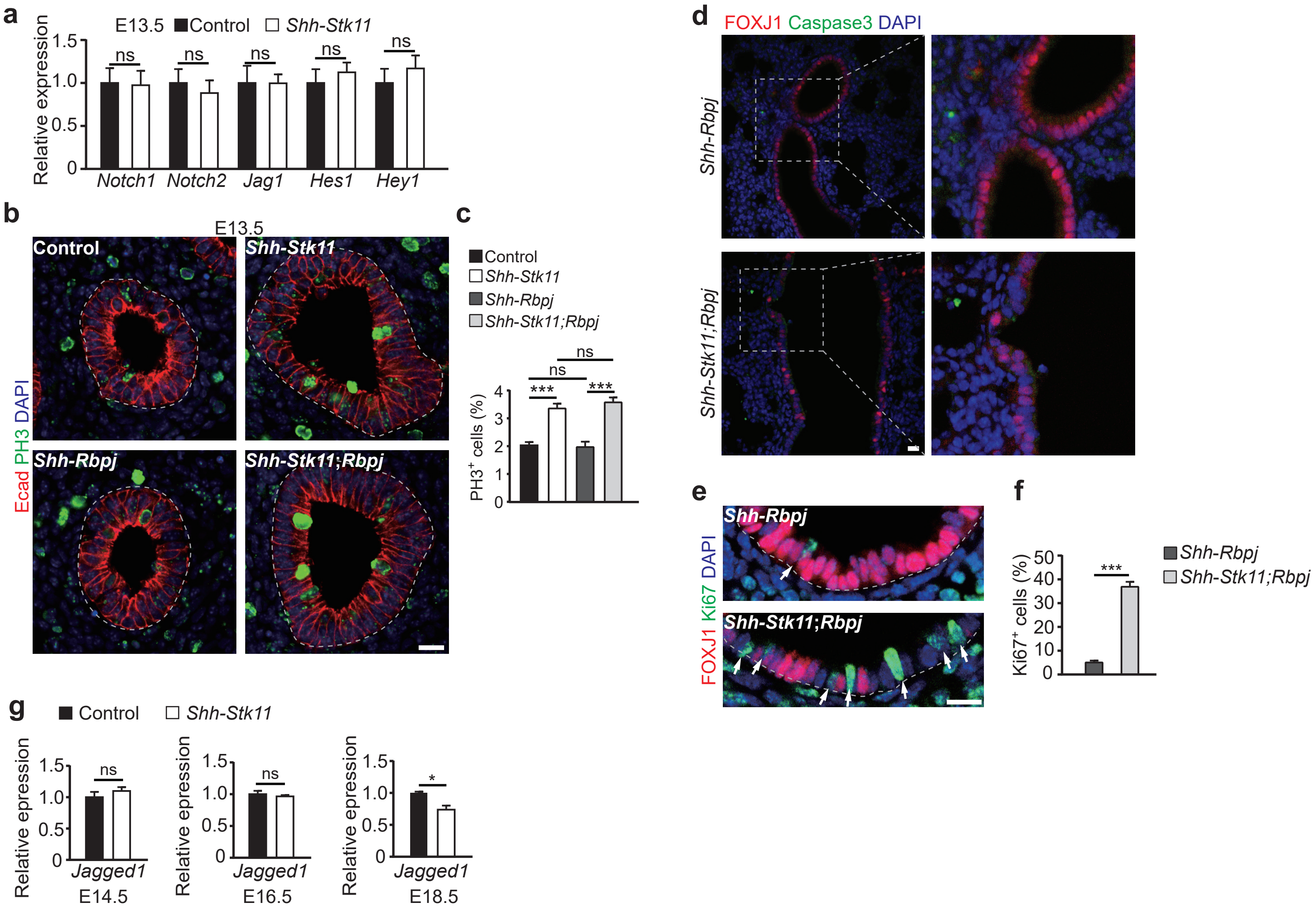
a
E13.5
Control
Shh-Stk11
Relative expression
Notch1
Notch2
Jag1
Hes1
Hey1
ns
b
E13.5
Control
Shh-Stk11
Shh-Rbpj
Shh-Stk11;Rbpj
Ecad PH3 DAPI
c
Control
Shh-Stk11
Shh-Rbpj
Shh-Stk11;Rbpj
PH3+ cells (%)
d
FOXJ1 Caspase3 DAPI
Shh-Rbpj
Shh-Stk11;Rbpj
e
FOXJ1 Ki67 DAPI
Shh-Rbpj
Shh-Stk11;Rbpj
f
Ki67+ cells (%)
Shh-Rbpj
Shh-Stk11;Rbpj
g
Control
Shh-Stk11
Relative epression
Jagged1
E14.5
E16.5
E18.5

**Figure S7.** The function of STK11 in regulating ciliated cell differentiation is independently of Notch signaling. **a** The mRNA expression level of Notch signaling pathway genes in the E13.5 Control lungs and *Shh-Stk11*$^{F/F}$ lungs. **b** Immunofluorescence staining with antibodies against PH3 and Ecad in E13.5 Control lungs, *Shh-Stk11*$^{F/F}$ lungs, *Shh-Rbpj*$^{F/F}$ lungs, and *Shh-Stk11*$^{F/F}$; *Rbpj*$^{F/F}$ lungs. **c** The proportion of PH3$^{+}$ airway epithelial cells in the intrapulmonary airways of E13.5 lungs (Control, n=4; *Shh-Stk11*$^{F/F}$, n=3; *Shh-Rbpj*$^{F/F}$, n=4; *Shh-Stk11*$^{F/F}$; *Rbpj*$^{F/F}$, n=3). **d** Immunofluorescence staining with antibodies against Caspase3 and FOXJ1 in E16.5 *Shh-Rbpj*$^{F/F}$ lungs and *Shh-Stk11*$^{F/F}$; *Rbpj*$^{F/F}$ lungs. **e** Immunofluorescence staining with antibodies against FOXJ1 and Ki67 in E16.5 lungs. White arrows indicate the FOXJ$^{-}$Ki67$^{+}$ cells. **f** The proportion of Ki67$^{+}$ cells in the intrapulmonary airways of E16.5 lungs (*Shh-Rbpj*$^{F/F}$, n=4; *Shh-Stk11*$^{F/F}$; *Rbpj*$^{F/F}$, n=6). **g** The mRNA level of *Jagged1* in the epithelia of Control and *Shh-Stk11*$^{F/F}$ lungs at E14.5, E16.5, and E18.5. ns, not significant, *, *P*<0.05; ***, *P*<0.001. Data shown in the graphs are means ± SEM. Student's *t*-test. Scale bars: **b**, **d**, **e**: 11 μm.

**Table S1. Sequences of DNA primers used for RT-qPCR**

| Gene | Forward sequence (5'-3') | Reverse sequence (5'-3') |
|---|---|---|
| *Gapdh* | CATCACTGCCACCCAGAAGACTG | ATGCCAGTGAGCTTCCCGTTCAG |
| *Myb* | GAAAGTGCCTCACCAGCAAGGT | CGAGCTTTCATGGTTGCTGGAAG |
| *Mcidas* | GTGGAAGTCCTTTTCGGGATGC | TAGGAGACGCTTCGGTTCCGAG |
| *Foxj1* | CTCCTATGCCACTCTCATCTGC | GACAGGTTGTGGCGGATGGAAT |
| *Scgb1a1* | GGTTATGTGGCATCCCTGAAGC | GCTTACACAGAGGACTTGTTAGG |
| *Muc5b* | CTGAAGACCTGTCGGAACCCAA | GCCACACACTTCATCTGGTCCT |
| *Ranbp2* | ATGGTTGCTGGCGAAGTGCTGA | ATCTTCTGCCCATCGAGGTGGT |
| *Smc5* | TGCGTGATACGAACAGGCACCT | GCAAGTATCCTGTTCCATCAGCC |
| *Mki67* | GAGGAGAAACGCCAACCAAGAG | TTTGTCCTCGGTGGCGTTATCC |
| *Sgo2a* | ACCACAGGACACAGAAGTCGAC | GTTGAGAGGGACGCACTGTCTT |
| *Taf1* | ATCCTACAGGCTGTGGTGAAGG | TTCAGAGACAGGCGACGAAGGT |
| *Ccnd2* | GCAGAAGGACATCCAACCGTAC | ACTCCAGCCAAGAAACGGTCCA |
| *Cenpe* | AGGATCATGCCACCGAGAAGAC | GCTGTGTCTCTTGGAGTTTCTGG |
| *Prkaa1* | GGTGTACGGAAGGCAAAATGGC | CAGGATTCTTCCTTCGTACACGC |
| *Prkaa2* | CTGAAGCCAGAGAATGTGCTGC | GAGATGACCTCAGGTGCTGCAT |
| *Mark1* | TGAAGGTCCAGCGGAGCATCTC | CCTCTTGGTGTATGAGACGGCA |
| *Mark2* | GAACCTCTCCCTGACTACAAGG | GGAGCAGATAGGTAGCCATCAC |
| *Mark3* | CAGTCTCCTCATCACAAAGGGC | TGCTGGTCTGACTCCTCTTTGG |
| *Mark4* | CTGCGGAGATTTCTGGTGCTGA | TCCGTGTATGGCTTCAGCTCCT |
| *Sik1* | GCGACTACAACGAACAGGTGCT | GTAGGAGGTAGTAAATGGCGGC |
| *Sik2* | GCTGCCTTTATGGAGGAAGAGTG | GAGGTCTCCATCATACTGCTGG |
| *Sik3* | AACGGCAGCTAGGACAACAGTC | GTGTTGGAGTCCTTGTAGGTGG |
| *Brsk1* | GACGTTCTGGAAAGCATGGCGT | GGTCTTCACAGCTAGGATACCG |
| *Brsk2* | CGGAAAGAACGGTATCCAAGCC | CATCTGTCACACTGAGCACTTCC |
| *Nuak1* | CAGATCAGCAGCGGAGAGTACC | CAATGGTTGGCGATGTCCTCGA |
| *Nuak2* | CTGGTGAAGCAAATCAGTAACGG | CCACCAATGACTGGCTACATCC |
| *Notch1* | GCTGCCTCTTTGATGGCTTCGA | CACATTCGGCACTGTTACAGCC |
| *Notch2* | CCACCTGCAATGACTTCATCGG | TCGATGCAGGTGCCTCCATTCT |
| *Jag1* | TGCGTGGTCAATGGAGACTCCT | TCGCACCGATACCAGTTGTCTC |
| *Hes1* | GGAAATGACTGTGAAGCACCTCC | GAAGCGGGTCACCTCGTTCATG |
| *Hey1* | CCAACGACATCGTCCCAGGTTT | CTGCTTCTCAAAGGCACTGGGT |
